# Fine-tuning of Epithelial EGFR signals Supports Coordinated Mammary Gland Development

**DOI:** 10.1101/2020.09.10.291872

**Authors:** Alexandr Samocha, Hanna M. Doh, Vaishnavi Sitarama, Quy H. Nguyen, Oghenekevwe Gbenedio, Joshua D. Rudolf, Walter L. Eckalbar, Andrea J. Barczak, Yi Miao, K. Christopher Garcia, Devon Lawson, Zena Werb, Kai Kessenbrock, Philippe Depeille, Jeroen P. Roose

## Abstract

During puberty, robust morphogenesis occurs in the mammary gland; stem- and progenitor-cells develop into mature basal- and luminal-cells to form the ductal tree. The receptor signals that govern this process in mammary epithelial cells (MECs) are incompletely understood. The EGFR has been implicated and here we focused on EGFR’s downstream pathway component Rasgrp1. We find that Rasgrp1 dampens EGF-triggered signals in MECs. Biochemically and *in vitro*, *Rasgrp1* perturbation results in increased EGFR-Ras-PI3K-AKT and mTORC1-S6 kinase signals, increased EGF-induced proliferation, and aberrant branching-capacity in 3D cultures. However, *in vivo, Rasgrp1* perturbation results in delayed ductal tree maturation with shortened branches and reduced cellularity. *Rasgrp1*-deficient MEC organoids revealed lower frequencies of basal cells, the compartment that incorporates stem cells. Molecularly, EGF effectively counteracts Wnt signal-driven stem cell gene signature in organoids. Collectively, these studies demonstrate the need for fine-tuning of EGFR signals to properly instruct mammary epithelium during puberty.

## Introduction

The mammary gland is a dynamic organ comprised of a network of bi-layered ductal epithelial structures that develops predominantly during puberty. In the ductal phase, during puberty, robust proliferation and branching morphogenesis need to be coordinated to assure development of the mature “ductal tree” from its rudimentary, embryonic tree (for reviews see (Hinck & Silberstein, 2005; Huebner & Ewald, 2014; Sternlicht & Sunnarborg, 2008; Wiseman & Werb, 2002)). The epithelial lineages that form these ducts arise out of terminal end buds, or TEBs, which consist of a mass of inner body cells surrounded by cap cells. Mammary stem cells (MaSCs) and mammary epithelial progenitor cells in this inner mass make up a heterogeneous cell population (Visvader & Stingl, 2014) that gives rise to polarized mammary ducts with a central luminal epithelial cell layer and a basally located myoepithelial layer located at the basement membrane (Huebner & Ewald, 2014; Wiseman & Werb, 2002).

The receptor signals that that govern the coordination of the outgrowth of epithelium during puberty in the mammary gland are still being defined. The EGFR (Epidermal Growth Factor Receptor) family consists of EGFR (ErbB1/Her1), ErbB2/Her2, ErbB3/Her3, and ErbB4/Her4 with ligands that include EGF, transforming growth factor-alpha (TGFα), and amphiregulin (AREG) (Hynes & Watson, 2010; Samocha, Doh, Kessenbrock, & Roose, 2019). Elegant mixed tissue- and transplantation-approaches into cleared fat pad defined an essential role for EGFR signaling in stromal cells to promote ductal development (Sebastian et al., 1998; Sternlicht et al., 2005; Wiesen, Young, Werb, & Cunha, 1999).

Surprisingly, the exact role that the EGFR plays in mammary epithelial cells (MECs) has remained rather elusive. Genetic mouse models suggest that the EGFR signals in the mammary epithelium; deletion of *ErbB2* results in impaired ductal outgrowth (Andrechek, White, & Muller, 2005; Jackson-Fisher et al., 2004), *ErbB3* deficiency leads to mammary outgrowth defects with smaller TEBs and increased branch density (Jackson-Fisher et al., 2008), and *ErbB4* deletion impacts the mammary gland during lactation but not during puberty (Tidcombe et al., 2003). Furthermore, expression of a dominant negative form of the EGFR from the MMTV promoter inhibits pubertal mammary duct development with reduced proliferation in the ducts (Xie, Paterson, Chin, Nabell, & Kudlow, 1997). The strength of EGFR signal also appears to matter; *Waved-2* mice carry a point mutation near the kinase domain of the EGFR, which reduces kinase activity (Luetteke et al., 1994). *Waved-2* females display impaired mammary development with reduced branching and decreased ductal invasion into the fat pad during puberty (Fowler et al., 1995; Sebastian et al., 1998).

Growth factor receptors, like the EGFR, signal through Ras guanine nucleotide exchange factors (RasGEFs) to activate the small GTPase Ras and kinase pathways that lie downstream of activated Ras, such as PI3K (Phosphatidylinositol-3 kinase) (Vigil, Cherfils, Rossman, & Der, 2010). The RasGEFs SOS1 and RasGRP1 are structurally very different (Iwig et al., 2013; Margarit et al., 2003; Vercoulen et al., 2017) and generate distinct Ras-kinase signals in lymphocytes (Daley et al., 2013; Das et al., 2009; Jun, Rubio, & Roose, 2013; Jun, Yang, Chen, Chakraborty, & Roose, 2013; Kortum, Rouquette-Jazdanian, & Samelson, 2013; Ksionda, Limnander, & Roose, 2013; Roose, Mollenauer, Ho, Kurosaki, & Weiss, 2007). We previously established that Rasgrp1 primes the activity of SOS1 in lymphocytes (Das et al., 2009) but that Rasgrp1 dampens EGFR-SOS1 signals in intestinal epithelial cells (Depeille et al., 2015). Here we characterized the role of EGFR signaling in maturation of the mammary gland during puberty. Having noted Rasgrp1 expression in the mammary gland, we capitalized on two genetic *Rasgrp1* mouse models to understand the impact of strength of EGFR signals on mammary development during puberty. We find that precisely tuned epithelial EGFR signals are essential for regulated MEC proliferation in maturing ducts, ductal tree maturation, and production of differentiated basal and luminal cells. *In vitro* organoid assays provide mechanistic insights and reveal that EGFR signals counteract a Wnt- and Rspondin-driven gene signature in of mammary epithelium cells.

## Results

### Impaired Rasgrp1 function results in increased EGFR-effector kinase signaling in mammary epithelial cells

SOS1 and RasGRP1 both activate Ras but display different molecular regulation (Iwig et al., 2013; Margarit et al., 2003; Vercoulen et al., 2017). SOS1 is ubiquitously expressed and is allosterically activated by RasGTP (Sondermann et al., 2004), which generates a positive feedback loop and triggers digital Ras-ERK signals (Das et al., 2009). By contrast, Rasgrp1 is an analog Ras activator (Das et al., 2009; Iwig et al., 2013; Vercoulen et al., 2017). Initially, Rasgrp1 was thought to be a T cell-specific Ras activator (Dower et al., 2000). However, since the report of a T cell developmental block in *Rasgrp1* deficient mice, many more expression sites for Rasgrp1 have been reported (Reviewed in (Ksionda et al., 2013)). In the intestine, Rasgrp1 dampens EGFR-SOS1 signals and downstream RasGTP-ERK signals (**Figure 1A**) (Depeille et al., 2015). Deletion of Rasgrp1 leads to increased proliferation of intestinal progenitor cells and increased numbers of differentiated Goblet cells (Depeille et al., 2015), as well as to accelerated tumor growth in mouse models of colorectal cancer (Depeille et al., 2015; Gbenedio et al., 2019).

**Figure 1.**
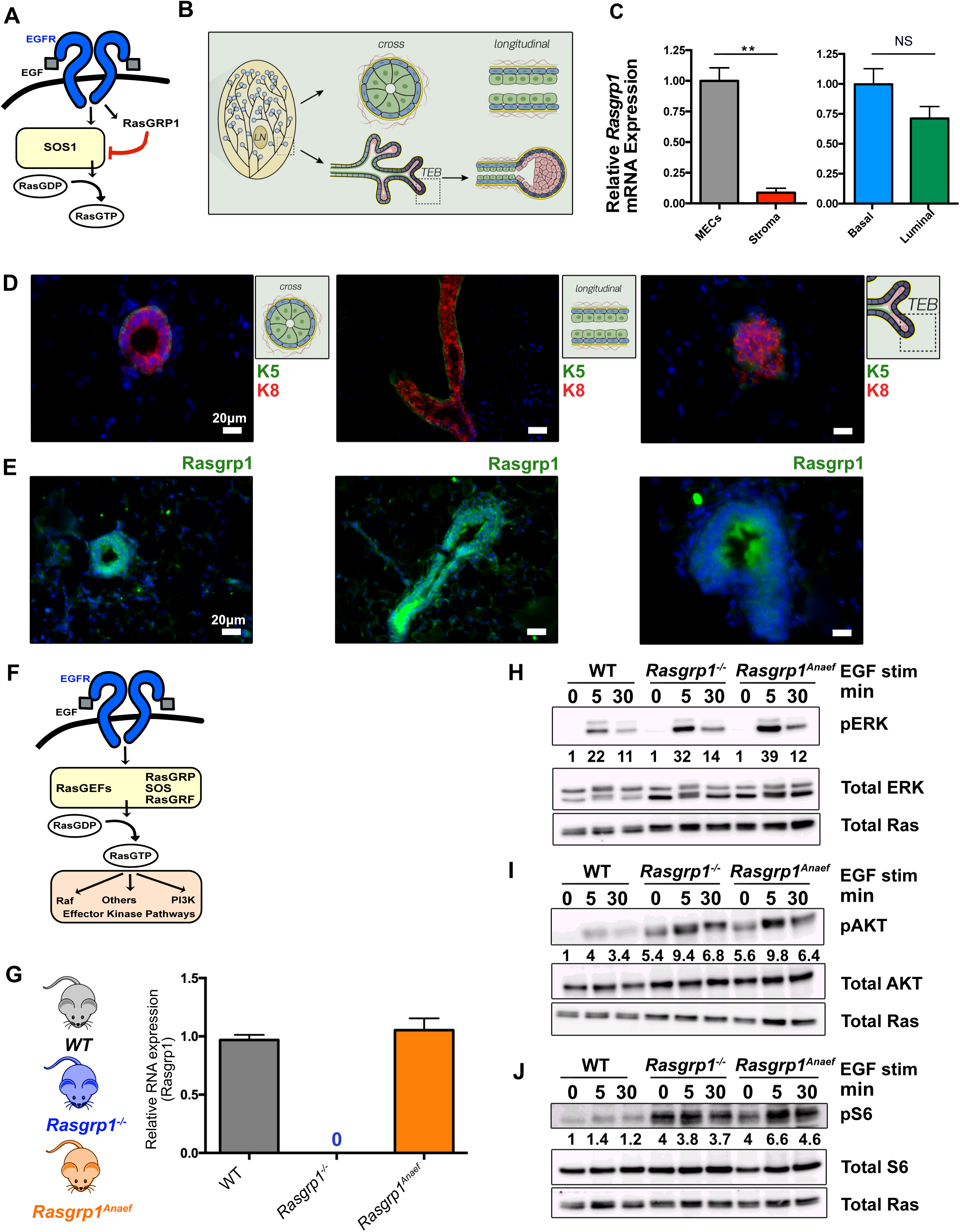
Rasgrp1 is expressed in the mammary epithelium during pubertal development and functions as a dampener of EGFR signals. (A) Cartoon of Rasgrp1 as a brake on the EGFR-SOS1-Ras pathway in the intestine. (B) Cartoon depicting the architecture of the mammary ductal network and duct and TEB (terminal end bud) make-up. (B) *Rasgrp1* mRNA expression in MECs. RNA was isolated from total MECs, sorted basal and luminal cells, and the surrounding stromal cells. Taqman RT-PCR was performed for *Rasgrp1* transcripts. Expression normalized to βActin RNA levels. 2^-ddct^ was calculated, with total MECs set to 1. t tests were performed for significance in pairwise comparisons. n = 3 for MECs and stroma, n = 2 for basal and luminal. (**p < 0.005, NS = not significant). (D, E) Immunofluorescence (IF) staining for mammary epithelial cell (MEC) markers in a wild-type (WT) C57BL/6 mammary gland. Cytokeratin-5 (K5, green) marks basal cells and cytokeratin-8 (K8, red) marks luminal cells. Cartoons depict positions in epithelial ductal tree and possible sections. (E) IF for Rasgrp1 expression (green) in luminal and basal MECs in 8-week-old WT mammary glands. DAPI (blue) counterstain marks cell nuclei. Representative examples are shown. Scale bar represents 20μm. (F) Cartoon of EGFR-RasGEF-Ras-kinase pathways. (G) *Rasgrp1* mRNA expression in MECs. RNA from isolated MECs was subjected to Taqman RT-PCR for *Rasgrp1* transcripts. Expression was normalized to βActin RNA levels. n = 4 per genotype. (H-J) EGF-induced ERK, AKT, and S6 kinase signals in wild type, *Rasgrp1^-/-^,* and *Rasgrp1^Anaef^* MECs. Western blot detection of phosphorylated ERK-, AKT-, and S6-proteins after 0, 5, and 30 minutes of EGF stimulation. Total ERK, AKT and S6 is consistent between samples. Total Ras serves as additional loading control. Ratiometric densitometry quantification of P-ERK/ERK, P-AKT/AKT, and P-S6/S6 is indicated below the blots with WT 0 stimulation arbitrarily set at 1.0. Blots in 1H-1J are representative examples of three independent experiments.

Various open access gene expression databases suggested Rasgrp1 expression in breast tissue and lineages (data not shown). We sorted mammary epithelial cell (MEC) subpopulations by flow cytometry and performed Taqman analysis for *Rasgrp1* mRNA expression (**Figure 1B** **and Supplemental Figures S1A and S1B**). Rasgrp1 was expressed in MECs but displayed negligible expression in the stroma (**Figure 1C**). CD31^-^/CD45^-^/Ter119^-^ “lineage-negative” cells (excluding endothelial cells, lymphocytes, and erythrocytes) can be divided into basal and luminal cells via analysis of EpCAM and CD49f expression (Kessenbrock et al., 2017; Stingl et al., 2006). *Rasgrp1* expression was detected in both basal and luminal cells (**Figure 1C**). Using the topology and make-up of mammary ducts and TEBs (**Figure 1B**) and dual staining for the basal cell marker Cytokeratin-5 and luminal Cytokeratin-8 marker for orientation (**Figure 1D**), we confirmed Rasgrp1 protein expression in both ducts and TEBs (**Figure 1E**).

Having established Rasgrp1 expression in epithelial cells of the mammary gland, we next explored two genetic *Rasgrp1* mouse models with the goal to better understand EGFR-Ras-kinase signaling in mammary gland epithelial cells and development (**Figure 1F**). We employed the *Rasgrp1*-deficient mouse model (Dower et al., 2000), but also a *Rasgrp1^Anaef^* model with an arginine to glycine mutation at position 519 (Arg 519 Gly) in *Rasgrp1*. T cell development is largely intact in *Rasgrp1^Anaef^* mice, but mice show autoimmune features around 20 weeks of ager (Daley et al., 2013). In the current study we used *Rasgrp1^Anaef^* mice at 11 weeks or younger. This *Rasgrp1^Anaef^* mutation has a mild Rasgrp1 autoinhibitory defect in resting cells but causes severely hypomorphic, receptor-induced Rasgrp1-Ras-kinase signals in stimulated cells (Daley et al., 2013). All mice were on a C57BL/6 background, which have somewhat slower pubertal development than FVB mice (**Supplemental Figure S1C**).

We isolated total MECs from wild type (WT), *Rasgrp1^Anaef^*, or *Rasgrp1*-deficient mice, and ran Taqman analyses for *Rasgrp1* mRNA levels demonstrating that *Rasgrp1* is roughly equally expressed in MECs from wild-type and *Rasgrp1^Anaef^*, and *Rasgrp1* is absent in *Rasgrp1^-/-^* MECs as expected (**Figure 1G**). Western blot analysis of isolated, total MECs confirmed expression of Rasgrp1 in both wild type and *Rasgrp1^Anaef^* MECs (**Supplemental Figure S1D**). *Rasgrp1^Anaef^* MECs have a slight reduction in total Rasgrp1 protein, as we have observed previously in T cells (Daley et al., 2013).

We used these MECs with *Rasgrp1* perturbations to test if Rasgrp1 functions as a primer or a brake on EGFR signaling in mammary epithelial cells. EGF stimulation of *Rasgrp1^-/-^* and *Rasgrp1^Anaef^* MECs resulted in modestly enhanced ERK phosphorylation (P-ERK), a downstream effector of the Ras-RAF-MEK kinase pathway, compared to the quantitated signal in EGF stimulated wild type MECs (**Figure 1H**). As downstream effector of PI3K signaling, AKT promotes protein synthesis, cell proliferation, cell metabolism and cell survival (Wickenden & Watson, 2010). We observed substantially elevated baseline AKT phosphorylation (P-AKT) in *Rasgrp1^-/-^* and *Rasgrp1^Anaef^* MECs that was further induced by EGF stimulation (**Figure 1I**). ERK and AKT can both connect to the ribosomal S6 signaling pathway, which plays roles in translation and metabolism. *Rasgrp1^-/-^* and *Rasgrp1^Anaef^* MECs also displayed robustly increased baseline levels of pS6 compared to wild type (**Figure 1J**). In sum, perturbations in *Rasgrp1* result in increased signals through ERK-, AKT-, and S6-effector kinase pathways in total MECs and Rasgrp1 functions as a dampener of EGFR-kinase signals in mammary epithelial cells, analogous to its inhibitory role in intestinal epithelial cells (Depeille et al., 2015; Gbenedio et al., 2019).

### Rasgrp1 perturbation leads to developmental delay in the ductal phase but increased branching in cleared fat pad assays

During puberty, bulbous TEBs form at the tips of ducts. Coordinated cell proliferation and migration drives invasion of TEBs into the fat pad, and branching occurs through TEB bifurcation or secondary side-branch sprouting (**Supplemental Figure S1C**) (Hinck & Silberstein, 2005; Huebner & Ewald, 2014; Sternlicht & Sunnarborg, 2008; Wiseman & Werb, 2002). To assess the impact of *Rasgrp1* perturbation on the ductal phase during puberty, we analyzed 5-, 7-, 9-, and 11- week-old C57BL/6 mice to capture pubertal development stages (**Supplemental Figure S1C**). *Rasgrp1^-/-^* or *Rasgrp1^Anaef^* females revealed an increase in TEB numbers at 7, 9, and 11 weeks (**Figures 2A and 2B and Supplemental Figure S2A**). The presence of TEBs in 11 week-old *Rasgrp1^-/-^* or *Rasgrp1^Anaef^* females is remarkable, as these structures have normally vanished by this age. The increased TEB numbers throughout puberty was accompanied by reduced lengthening of the ductal tree in *Rasgrp1^-/-^* or *Rasgrp1^Anaef^* females, which had a runted appearance (**Figures 2C and 2D and Supplemental Figure S2B**). We utilized FACS (Fluorescence-activated Cell Sorting) to more precisely enumerate and characterize MECs (Inman, Robertson, Mott, & Bissell, 2015). We first determined the absolute cellularity of lineage-negative cells that are also CD49f^+^/EpCAM^+^ in the mammary gland. Eight-week-old *Rasgrp1^-/-^* or *Rasgrp1^Anaef^* females demonstrated a significant reduction of total CD31^-^/CD45^-^/Ter119^-^ lineage-negative cells measured by quantitative FACS analysis with co-analyzed counting beads (**Figure 2E**). Capitalizing on analysis of EpCAM and CD49f expression (Kessenbrock et al., 2017) to further analyze subsets, we established that WT, *Rasgrp1^-/-^*, and *Rasgrp1^Anaef^* females all displayed luminal CD49f^medium^/EpCAM^high^ and basal CD49f^high^/EpCAM^medium^ cells (**Figure 2F**). Thus, there is no absolute impairment in development of either basal- or luminal-cell types in *Rasgrp1^-/-^* and *Rasgrp1^Anaef^* females.

**Figure 2.**
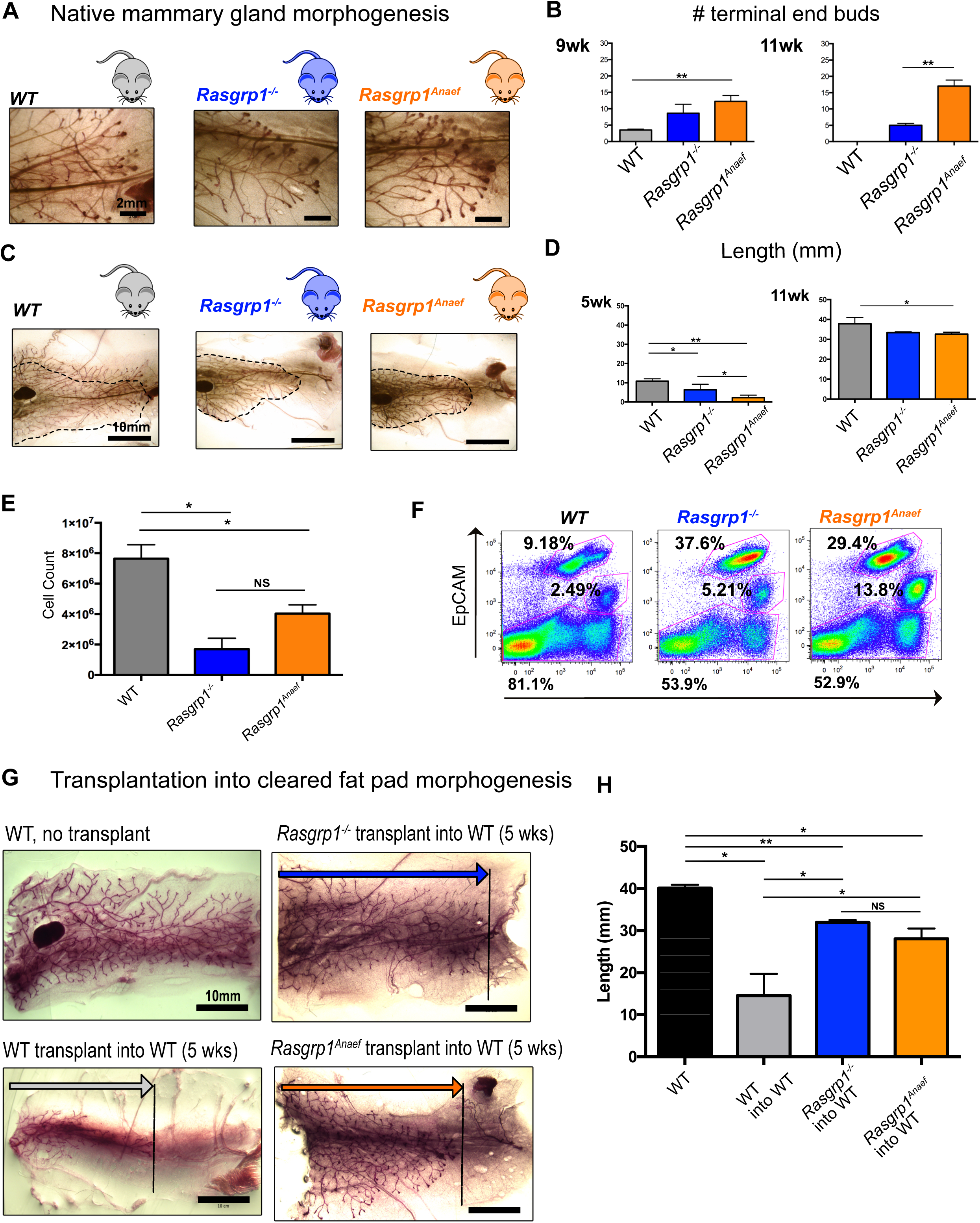
*Rasgrp1^-/-^* and *Rasgrp1^Anaef^* mice display delayed native mammary gland morphogenesis but enhanced features in cleared fat pad transplantations. (A, B) Mice with Rasgrp1 perturbations show elevated number of the proliferative epithelial structures, terminal end buds (TEBs). Representative images from 7-week-old, carmine-stained whole mounts. Scale bar, 2 mm. (B) Quantification of TEBs. *Rasgrp1^-/-^* and *Rasgrp1^Anaef^* mice show increased numbers of TEBs compared to their WT counterparts. Statistical significance determined by t test. (*p < 0.05, **p < 0.005; n=4). See also Supplemental Figure S2A. (C, D) Rasgrp1 perturbations result in decreased ductal elongation. 7-week-old mammary whole mounts from WT, *Rasgrp1^-/-^*, and *Rasgrp1^Anaef^* were carmine stained and imaged on a stereo dissection scope. Scale bar, 10mm. (D) Quantification of ductal elongation of 5-, and 11-week old mice. Length is measured from the nipple-proximal end of the lymph node to the furthest epithelial branch. At early pubertal development, *Rasgrp1^-/-^* and *Rasgrp1^Anaef^* have significantly shorter ductal elongation (*p < 0.05, **p < 0.005; n= 4). Statistical significance determined by t test. This impairment persists through 7, 9, and 11 weeks of age. See also Supplemental Figure S2B. (E) Quantitative FACS analysis was employed to count the total number of CD49f-EpCAM positive mammary epithelial cells in the indicated mouse models at 8 weeks of age. *p < 0.05, n= 4 per group. t tests performed for significance. (F) Representative EpCAM and CD49f FACS analyses of mammary epithelial cells gated on lineage negative live singlets. Each mouse model is a pool of two age-matched littermates. (G) Representative images on wild type, *Rasgrp1^-/-^,* and *Rasgrp1^Anaef^* mammary epithelial cells (MECs) transplanted into recipient cleared 5-week-old WT mammary fat pad. Ductal elongation was allowed for 6 weeks before images were acquired. Stereo dissection scope-capture of whole mounted and carmine stained mammary gland. Scale bar, 10mm. n=3, total mice transplanted per genotype. As a reference a non-transplanted mammary gland is shown. (H) Quantification of mammary transplant branch lengths. Wild-type (WT) non- transplanted glands are compared to WT, *Rasgrp1^-/-^* and *Rasgrp1*^Anaef^ transplants. n=3, total mice transplanted per group. Significance determined by t test. (*p < 0.05; **p < 0.005).

To further assess the developmental and proliferative potential of *Rasgrp1^-/-^* or *Rasgrp1^Anaef^* MECs, we transplanted isolated cells into cleared mammary fat pads of 5-week-old wild type recipient mice with the mammary gland at the contralateral side as an internal control (Kessenbrock et al., 2013; Lawson, Werb, Zong, & Goldstein, 2015) (**Supplemental Figure S2C**). Transplantation of wild type MECs into wild type recipients resulted in a developing ductal tree when analyzed 6 weeks post-transplant. Unexpectedly, transplantation of *Rasgrp1^-/-^* or *Rasgrp1^Anaef^* cells into a cleared fat revealed a more advanced branched tree (**Figure 5E** **and 5F**). Thus, *Rasgrp1^-/-^* and *Rasgrp1^Anaef^* MECs have intact intrinsic capacity to form a ductal tree when assessed in cleared fat pad assays and can give arise to both luminal and basal cell populations, yet, both *Rasgrp1^-/-^* and *Rasgrp1^Anaef^* perturbations leads to reduced cellularity in native mammary gland morphogenesis, reduced lengthening of the ductal tree, and persistent presence of TEBs late in puberty.

### Rasgrp1 suppresses proliferation and AKT signals

To mechanistically understand the effects of *Rasgrp1* perturbation in mammary epithelium, we digested the mammary gland with enzymes into single cells and subsequently plated these to probe MEC proliferative capacity in sphere-forming assays (**Figure 3A**). Using this assay, we observed increased colony formation for *Rasgrp1^-/-^* or *Rasgrp1^Anaef^* compared to WT MECs (**Figure 3B**), which was dependent on EGFR kinase signaling (**Supplemental Figure S3A**). Immunohistochemistry on the spheres revealed that the larger and more numerous *Rasgrp1^-/-^* or *Rasgrp1^Anaef^* colonies displayed high levels of P-AKT (**Figure 3C**), which was abrogated by the inclusion of the EGFR inhibitors (**Supplemental Figure S3B**). The *Rasgrp1^-/-^* and *Rasgrp1^Anaef^* colonies also displayed high levels of the proliferation marker Ki67 (**Figure 3D**).

**Figure 3.**
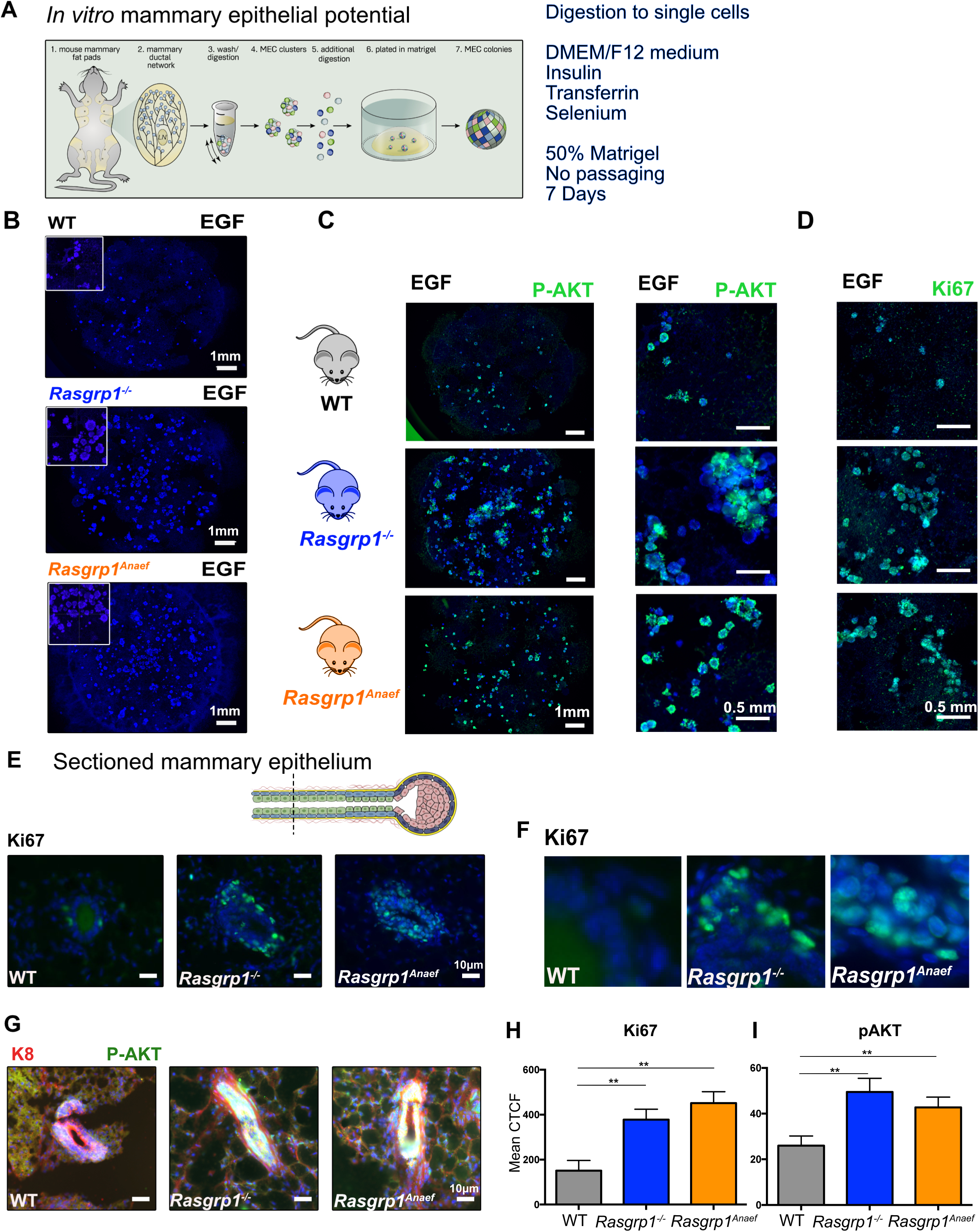
Progenitor cell proliferation and AKT signals are increased in Rasgrp1^-/-^ and Rasgrp1^Anaef^ females. (A) Cartoon representation of mammary epithelial cell colony forming assay. Single cells are placed into Matrigel pellet and followed for 7 days to assay colony number and size. (B) *Rasgrp1^-/^*^-^ and *Rasgrp1^Anaef^* single MECs form more numerous and larger clusters in response to EGF compared to WT. In each Matrigel pellet, 2500 cells were loaded. Addition of EGFR inhibitors (EGFRi) blocks the growth of all colonies. Hoechst 33342 is used as nuclear counterstain. Scale bar, 1 mm. Zoom in images (top left). (C) AKT phosphorylation of colony formation assay on the three indicated, plated single MEC populations. Scale bar, 1 mm. (Right) Higher magnification of images with scale bar, 0.5 mm. (D) Ki67 staining of colony formation assay on the three single MEC populations plated to assess cell proliferation. Scale bar, 0.5 mm. (E, F) Ki67 immunofluorescence (green) on cross-sections of terminus proximal mammary ducts from 9-week-old WT, *Rasgrp1^-/-^*, and *Rasgrp1^Anaef^* mice. DAPI (blue) counterstain marks cell nuclei. Representative examples are shown. Scale bar, 10 μm. (F) Higher magnification of Figure 2E. (G) AKT phosphorylation (green) by IHC on cross-sections of terminus proximal mammary ducts from 9-week-old WT, *Rasgrp1^-/-^*, and *Rasgrp1^Anaef^* mice. K8 (luminal) stained in red. DAPI (blue) counterstain marks cell nuclei. Exposure was equal in all three images and set to highlight that there is some P-AKT in WT. Representative examples are shown. Scale bar, 10 μm. (H) Quantification of the mean corrected total cell fluorescence (CTFC) for Ki67 IF. n = 4 for all groups. Significance determined by t test. (** p < 0.005). (I) Quantification of the mean CTCF for pAKT IF. n = 4 for all groups. Significance determined by t test. (** p < 0.005).

The Eph4 mammary epithelial cell line is devoid of Rasgrp1 expression (**Supplemental Figure S3C**), providing an “empty vessel/add-back” system (**Supplemental Figure S3D**). Stimulation of Eph4 cells with EGF resulted in robust outgrowth of colonies and colony numbers were reduced when EGFR inhibitors were added, thus, this system is responsive to EGFR signals (**Supplemental Figure S3E**). Retroviral expression of a wild type version of Rasgrp1 in Eph4 cells suppressed colony numbers, but expression of a catalytically inactive form of Rasgrp1 with an Arg_271_ to Glu substitution (R271E) did not (**Supplemental Figure S3F**). Thus, the catalytic activity of Rasgrp1 is required to suppress EGFR-driven proliferation (**Supplemental Figure S3G**).

We next investigated proliferation and AKT signals in sectioned ducts from our mouse models. At 9 weeks, when we observed increased TEB numbers and reduced length of the branched network, *Rasgrp1^-/-^* or *Rasgrp1^Anaef^* females display continued proliferation at duct locations that are supposed to contain fully differentiated cells, as evidenced by the high number of Ki67-positive, proliferative cells. By contrast, proliferation had decreased at that age in wild type females (**Figures 3E, 3F, and 3H**). Immunohistochemistry for P-AKT revealed increased levels in ducts from 9 week-old *Rasgrp1^-/-^* or *Rasgrp1^Anaef^* females (**Figure 2G** **and 2I)**. These observations agree with reports that AKT signals can promote proliferation in MECs in conjunction with some level of ERK signaling (Worster et al., 2012).

### Rasgrp1 perturbation elicits gain-of-function branching effects from EGF signals

Three-dimensional (3D) cultures of mammary epithelium have provided important insights in the complex biology of this organ (Mroue & Bissell, 2013). Digestion of the mammary gland to epithelial clusters (*i.e.* tissue chunks, not single cells) and plating in Matrigel allows for 3D *in vitro* studies on growth factors and cell biology (**Figure 4A**). Both WT and *Rasgrp1^-/-^* MEC displayed stereotypical organoid formation after 5 days when cultured in DMEM/F12 medium supplemented with Insulin-Transferrin-Selenium and 2.5 nM EGF as extrinsically added growth factor (Mroue & Bissell, 2013) (**Figure 4B**) and both genotypes yielded luminal CD49f^medium^/EpCAM^high^ as well as basal CD49f^high^/EpCAM^medium^ cell populations (**Figure 4C**).

**Figure 4.**
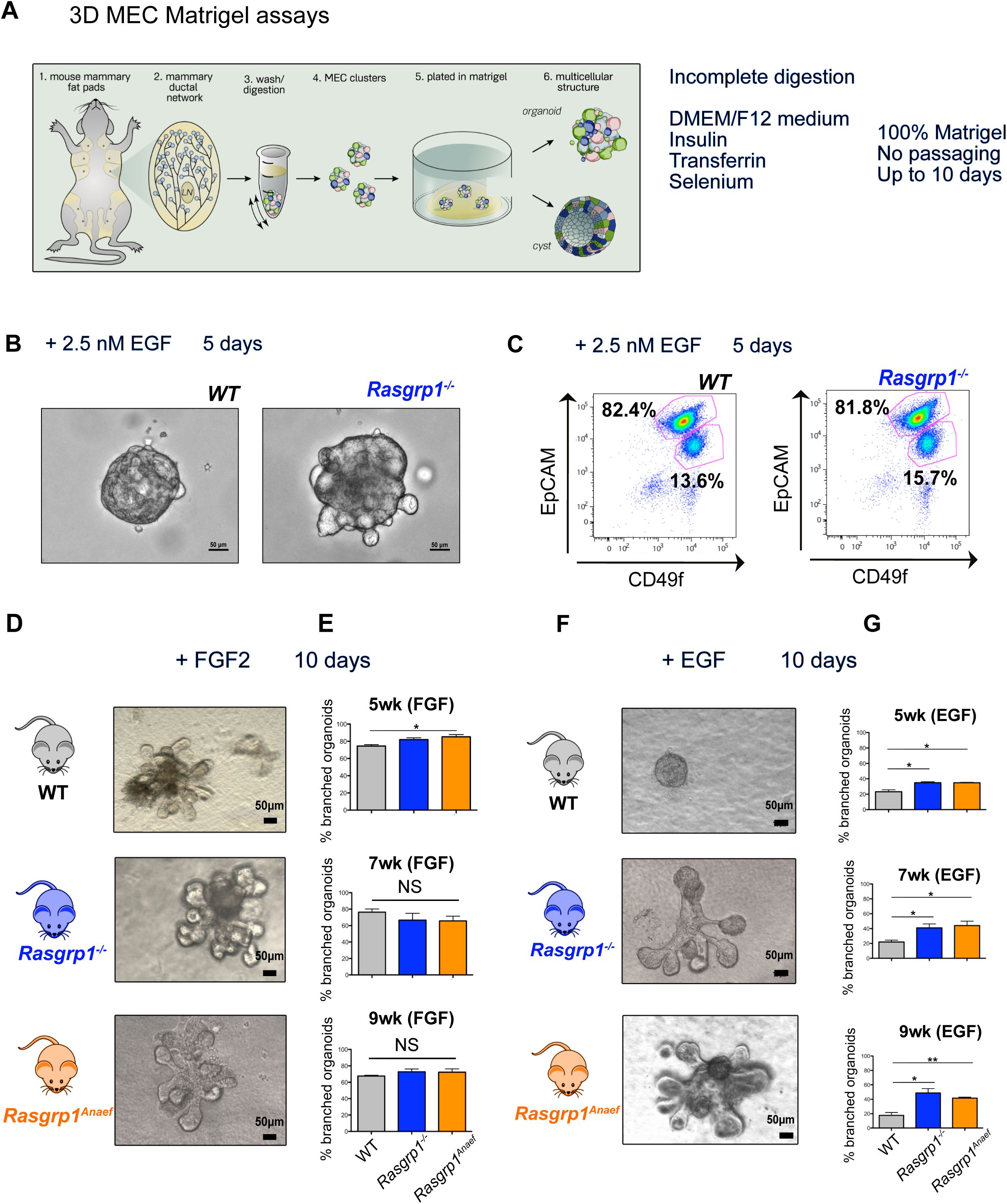
Perturbations to Rasgrp1 results in a gain-of-function EGF- induced branching in 3D mammary cultures. (A) Digestion and culture conditions with cartoon representation of the 3D MEC Matrigel assays. Isolated mammary fat pads are enzymatically digested. Differential centrifugation separates mammary epithelial cell clusters, which grow into 3D structures in Matrigel. (B, C) WT and *Rasgrp1^-/^*^-^ -derived 3D cultures with 2.5 nM EGF in the media, analyzed by imaging and EpCAM and CD49f FACS. Representative results of three independent experiments. (D, E) 3D cultures derived from WT, *Rasgrp1^-/^*^-^, *Rasgrp1^Anaef^* mice all respond robustly to the branching agonist FGF2. Representative images are shown for FGF2-stimulated branched 3D cultures derived from 7-week-old mice, imaged at day 10. At 5 weeks, *Rasgrp1^Anaef^* 3D cultures respond more strongly to FGF2 stimulation (*p < 0.05), n=3 for each genotype, time point. Differences between all other groups are not significant (NS). Significance determined by t test. Scale bar, 50 μm. (F, G) *Rasgrp1^-/^*^-^ and *Rasgrp1^Anaef^*-derived 3D cultures have augmented branching in response to EGF stimulation (Test? *p < 0.05, ** p < 0.005, n=4 for each genotype and time point). Representative 3D culture images are shown for each group, derived from 7-week-old mice. Scale bar, 50 μm.

FGF2 is known to potently promote branching of MECs in these mammary organoids over the course of 10 days (Ewald, Brenot, Duong, Chan, & Werb, 2008; Mroue & Bissell, 2013). As expected, FGF2 stimulated robust branching in 60 to 80% of our 3D Matrigel cultures from all three genotypes with a slight increase in branching for *Rasgrp1^Anaef^* from 5-week-old females (**Figures 4D and 4E**). Normally, EGF only induces spherical growth without substantial branching (Pasic et al., 2011), but *Rasgrp1^-/-^* or *Rasgrp1^Anaef^* organoids demonstrated substantial branching when stimulated solely with EGF (**Figures 4D and 4E**). These branches start out as buds coming of the sphere that can be counted on day 6 (**Supplemental Figure S4A**). The increased branching in *Rasgrp1^-/-^* or *Rasgrp1^Anaef^* organoids was reduced when EGFR inhibitors were added after 3 days, demonstrating that this gain-of-function was dependent on EGFR signals (**Supplemental Figure 4A and S4B**).

In sum, *Rasgrp1^-/-^* or *Rasgrp1^Anaef^* females demonstrate delayed native mammary development *in vivo*, but increased EGF-induced proliferation, AKT signaling, and branching patterns in our *in vitro* assays. We hypothesized that the coordination of different growth factor cues is lost when Rasgrp1 function is perturbed. In the mammary gland, EGFR signals act in concert with other receptors signals (Inman et al., 2015; Visvader & Stingl, 2014)), such as Wnt ligands binding to Frizzled receptors to sustain stem cells, and we next investigated the interplay between EGF and Wnt to place the phenotype seen in *Rasgrp1^-/-^* and *Rasgrp1^Anaef^* females in broader context.

### Understanding interplay of growth factor cues with organoids

Development of mammary organoid models is an active area of research; inclusion of Wnt3a and Rspo2 (R-spondin 2) in growth media facilitated organoid formation and budding that grow out to structures with internal, polarized luminal cells and an outer network of elongated myoepithelial cells (Jamieson et al., 2017). Wnt signals support stem cell function in many organs and enable efficient generation of organoids in Matrigel *in vitro* (Clevers, 2016; Sato & Clevers, 2013). Binding of the ligand R-spondin to the receptor Lgr5 (Leu-rich repeat-containing receptor 5) sustains Wnt signals (de Lau et al., 2011). Design and generation of surrogate Wnt ligands (DKK Dickkopf, combined with Wnt, **Figure 5A**) used in tandem with R-spondin triggers Lgr5 and LRP/Frizzled receptors to mimic sustained, canonical Wnt signaling and enables effective growth of organoids (Janda et al., 2017). “Next generation surrogate” (NGS) Wnts are the next version and have proven highly effective to initiate and expand organoids from multiple different types of tissues, including kidney, colon, pancreas, hepatocyte, ovarian and breast (Yi Miao et al., In press). Using colon organoids we have performed careful tittering of the Wnt signals and as low as 0.1nM NGS Wnt sustained colon organoids (Yi Miao et al., In press). =

In order to better understand the impact of different growth factors on WT and *Rasgrp1^-/-^* mammary epithelial cells, we plated MECs in simplest possible DMEM/F12 medium with Insulin-Transferrin-Selenium as before and systematically assessed the impact of added NGS Wnt/Rspo2 (RSpondin2), EGF, or the combination of Wnt/Rspo2/EGF on generated organoids. Importantly, no conditioned media components or other factors that are challenging to standardize were used. Titrating NGS Wnt and Rspo2 revealed that 0.1nM and 8.3nM, respectively, sustained healthy growth while allowing for additional cell biological effects induced by 2.5nM EGF (Data not shown). Furthermore, note that we purposely used the most basic media here in order to focus on the LRP5/6/Frizzled, Lgr5, and EGFR signals (**Figure 5A**) and that we did not add additional components such as FGF7, FGF10 and Noggin, described for breast cancer to sustain indefinite organoid growth and make living biobanks (Sachs et al., 2018).

Morphologically, as seen before, *Rasgrp1^-/-^* MEC organoids revealed some budding when exposed solely to EGF combination, but both WT and *Rasgrp1^-/-^* MEC organoids demonstrated robust budding when exposed to the combination of EGF-, Wnt-, and R-spondin signals (“*ERW*”) (**Figure 5B**). Capitalizing on FACS, we established that cocktails containing NGS Wnt/Rspo2 (“*RW*”) and EGF/NGS Wnt/Rspo2 (“*ERW*”) were roughly twice as efficient to sustain basal CD49f^high^/EpCAM^medium^ cells compared to EGF alone (“*EGF*”) at day 5 (**Figure 5C**). This CD49f^high^/EpCAM^medium^ population is described to contain stem cells (Samocha et al., 2019; Shackleton et al., 2006; Stingl et al., 2006). Further culture until day 13 in these three defined growth factor cocktails revealed three aspects; Firstly, only NGS Wnt/Rspo2 (“*RW*”) efficiently sustains basal cells (CD49f^high^/EpCAM^medium^) over time. Second, *Rasgrp1^-/-^* MEC organoids displayed lower percentages of basal CD49f^high^/EpCAM^medium^ cells in all three cocktails as compared to WT MEC organoids (**Figure 5E**). Lastly, the cell composition within the day 13 organoids, based on the CD31^-^/CD45^-^/Ter119^-^ lineage-negative and EpCAM/CD49f FACS profiling, is remarkably similar for EGF and EGF/NGS Wnt/Rspo2 (“*ERW*”) but unique for NGS Wnt/Rspo2 (“*RW*”) (**Figure 5E**). An unique FACS profile (See 22.8% and 21.4% in **Figure 5E**) is exclusively sustained by NGS Wnt/Rspo2 (“*RW*”) and disappears when 2.5 nM EGF is added to Wnt signals, and we next investigated with RNAseq if EGFR signals override Wnt signals in these organoids.

### Molecular programs driven by EGF-, Wnt-, and Rspondin-signals

To understand the molecular programs driven by EGF-, Wnt-, and Rspondin-signals and combinatorial stimuli we performed gene expression analysis by total RNAseq on day 5 organoids. Unsupervised clustering of gene expression patterns revealed that growth factors drove overall gene expression patterns and overpowered differences coming from the WT or *Rasgrp1^-/-^* (KO) genotype (**Figure 6A, Supplemental Figure 5, and Supplemental table**). The RW and EGF conditions are furthest removed from each other. Whilst ERW lies in the middle, ERW is not simply a combination of RW and EGF (**Figure 6A**). Principal component (PC) analysis confirmed these unsupervised clustering observations; PC1 covers 44% percent of the variance, segregating RW away from EGF and ERW (**Figure 6B**). Addition of PC2 and PC3 provided further resolution in the visualization of the overall differences in gene expression.

**Figure 5.**
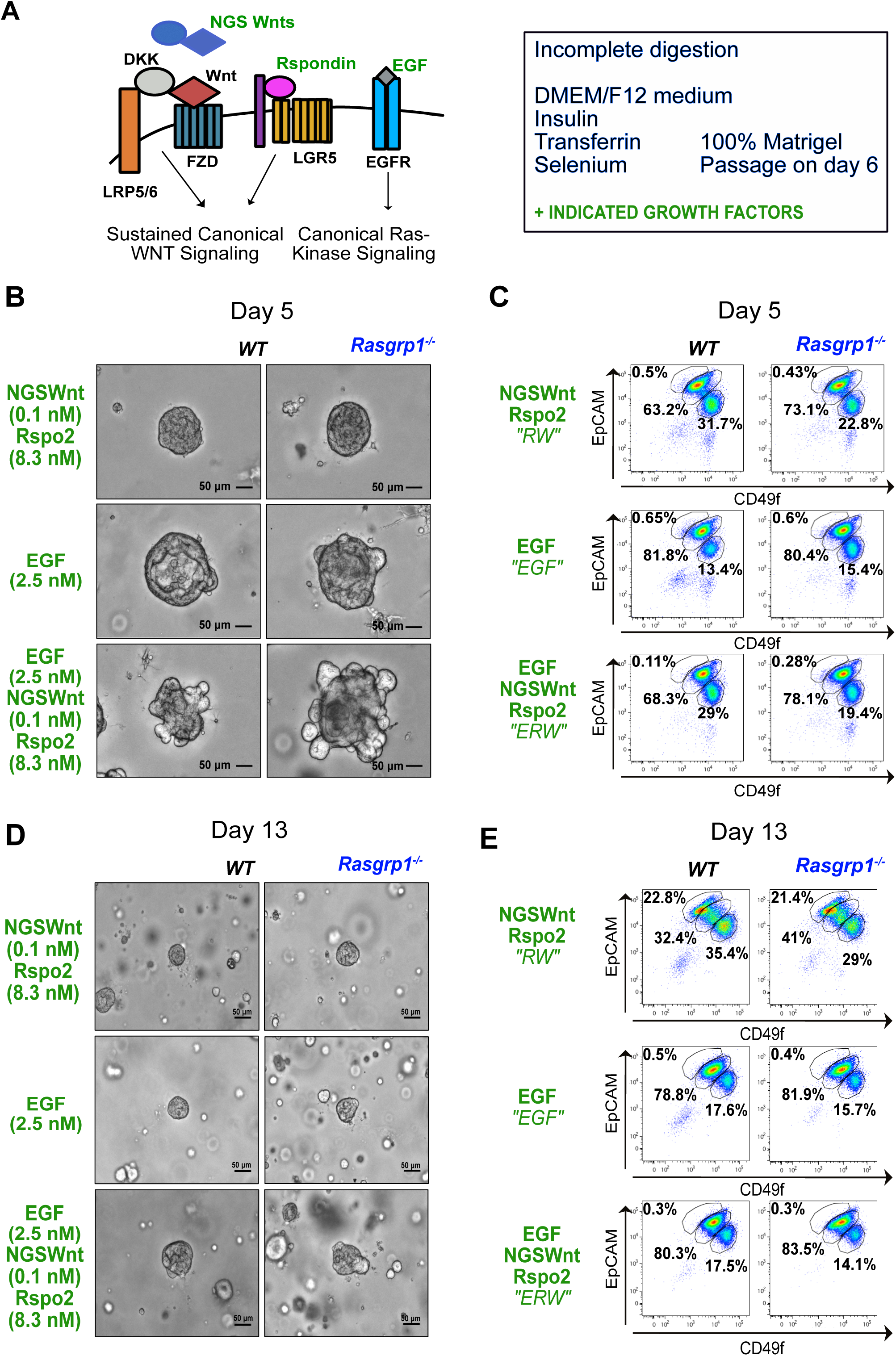
Organoids to understand coordinating signaling pathways. (A) Cartoon of Surrogate Wnt ligands (NGS Wnt), R-Spondin (Rspo), and EGF triggering canonical Wnt and Ras-Kinase pathways. Organoids are set-up in fully defined medium with precise growth factor concentrations and no conditioned media. (B) Morphology of wild type and *Rasgrp1^-/-^* mammary epithelial organoid assays in response to exposure to the indicated growth factors. Images are taken at day 5. Figures 5B-E are representative of three independent experiments. Each mouse model is a pool of two age-matched littermates. (C) Representative EpCAM and CD49f FACS analyses of mammary epithelial cells gated on lineage negative live singlets at day 5. The three gates indicate the relative percentage of cells with that marker phenotype. (D, E) As in Figures 5B and 5C, but images taken and FACS analysis performed at day 13. Organoids were passaged on day 6.

**Figure 6.**
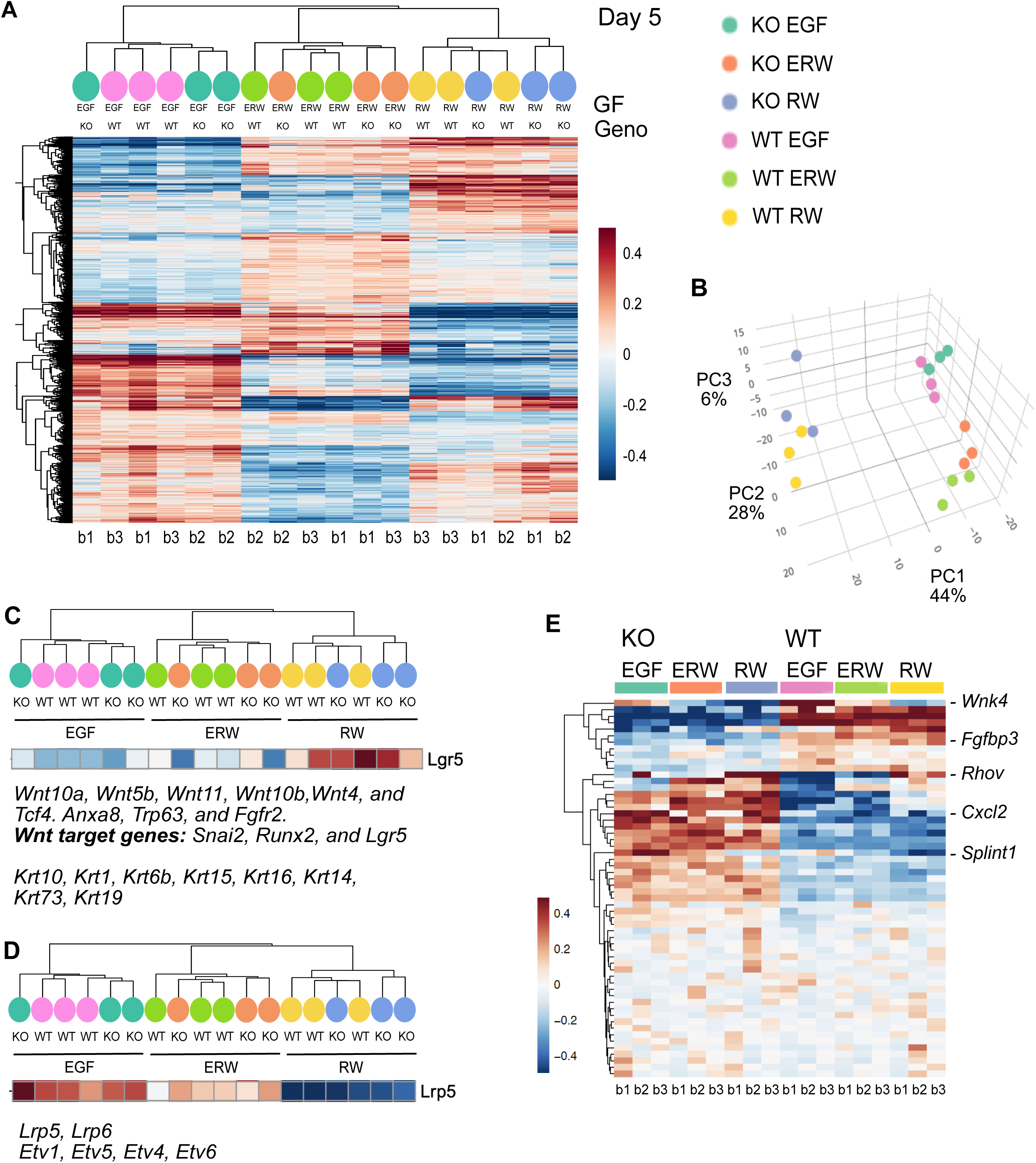
EGF suppresses Wnt/Rspondin-driven stem cell gene signatures. (A) Unsupervised clustering of total RNAseq data from wild type (WT) and *Rasgrp1^-/-^* (KO) mammary epithelial organoid assays at day 5. Each RNA sample is a pool of two age-matched littermates. The experiment was performed in three batches; b1-b3. The significance cut-off for Figure 6A was false discovery rate <0.05. No fold change thresholding was used to restrict genes in 6A. Genotype and RW, ERW, and EGF growth factors are indicated. Raw data are deposited on GEO NCBI. (B) Principle component analysis of the eighteen generated gene signatures. Colo-coding as in figure 6A. (C) Summary of gene expression that is maintained by RW but lost by addition of EGF. Examples are several Wnt pathway components, Wnt signal modifiers, and Wnt target genes such as *Lgr5*. For a complete list see Supplemental figure S5. (D) Analysis of genes that are expressed at low levels in RW but induced by addition of EGF. (E) Analysis of gene expression differences in wild type (WT) and *Rasgrp1^-/-^* (KO) mammary epithelial organoids, irrespective of growth factor input. Differentially expressed genes were selected on the basis of difference between KO (EGF) and WT (EGF) with a false discovery rate of <0.05. For a complete list with gene names see Supplemental figure S6.

Based on results in Figure 5, we hypothesized that RW medium may uniquely sustain stem cell gene signatures in our organoids and that the addition of EGF could steer away from such signatures. To explore this, we investigated a gene set that was expressed at high level in RW medium but showed reduced expression levels in ERW (and also in EGF) (**Figure 6C** **and Supplemental Figure S5**). In this gene cluster we can identify Wnt signaling pathway components that were selectively expressed at high levels in RW conditions: *Wnt10a, Wnt5b, Wnt11, Wnt10b, Wnt4*, and *Tcf4*. Thus, (a portion of) cells in the RW organoids make their own Wnt ligands and express the transcription factor Tcf4 that is known to associate with beta-catenin to turn on target genes in response to Wnt signals (van de Wetering et al., 2002). Modulators of Wnt signals, *Anxa8* (Anexin A8), *Trp63* (p63), and *Fgfr2* are also in this cluster. Anxa8 is expressed in c-kit-positive luminal progenitor cells in the mammary epithelium (Iglesias et al., 2015) and can enhance Wnt target gene expression (Lueck, Carr, Yu, Greenwood, & Moss, 2020). p63 is essential for proliferation of epithelial stem cells (Senoo, Pinto, Crum, & McKeon, 2007) and promotes beta-catenin signaling (Lee et al., 2014), and Fgfr2 can promote beta-catenin signaling as well (Krejci et al., 2012). In further support of active Wnt signaling in the RW organoids, we observed selective high expression of Wnt target genes *Snai2 (Slug)* (Vallin et al., 2001; Wu et al., 2012), *Runx2* (Ferrari et al., 2015; Gaur et al., 2005; Van Keymeulen et al., 2011), and the target *Lgr5* (Barker, Tan, & Clevers, 2013) that is the receptor for R-spondin (de Lau et al., 2011) (**Figure 6C**). We also noted the selectively high expression of different keratins some of which have been described on epithelial progenitors in the mammary gland (Stingl, Raouf, Emerman, & Eaves, 2005); *Krt10, Krt1, Krt6b, Krt15, Krt16, Krt73, Krt19*, as well as Krt14, which was used in mammary stem cell tracing studies (Van Keymeulen et al., 2011). In sum, the NGS Wnt/Rspo2 (“*RW*”) media results in a gene signature rich for Wnt signaling and addition of EGF to RW diverts from this signature.

It should be noted that there is also a cluster of genes that is selectively expressed at low levels in RW conditions and induced by EGF. Interesting examples here are the Frizzled co-receptors *LRP5* and *LRP6* and the transcription factors *Etv1, Etv5, Etv4*, and *Etv6* (**Figure 6D** **and Supplemental table**). These Etv factors belong to the PEA3 subfamily of Ets transcription factors, which are nuclear effectors of Ras-Kinase signaling involved in cell proliferation and differentiation (Wasylyk, Hagman, & Gutierrez-Hartmann, 1998). Lastly, whereas the growth factor combination clearly dominate the gene expression profiles, we also explored the specific differences between organoids from WT and *Rasgrp1^-/-^* MECs for completeness. We clustered genes that revealed the same differences between the genotypes in all culture conditions. These analyses resulted in a limited list of genes expressed at lower levels or higher levels in *Rasgrp1^-/-^* MECs that tie in with various cell biological processes (**Figure 6E** **and Supplemental Figure 6**).

## Discussion

Here we demonstrate that perturbations in *Rasgrp1*, either via deficiency in *Rasgrp1* or *Rasgrp1^Anaef^*, delay mammary development *in vivo*. The ductal tree is short, TEB numbers are aberrantly present late in puberty, and increased proliferation and AKT signaling is observed where MECs should be terminally differentiated and not dividing. Biochemically and *in vitro*, Rasgrp1 perturbation results in increased EGFR-Ras-PI3K-AKT and mTORC1-S6 kinase signals, and increased proliferative- and branching-capacity in response to EGF. These results demonstrate that EGFR signals in the mammary epithelium need to be fine-tuned. In support of that notion, we show that EGF is effective to negate NGS Wnt and Rspondin2-driven Wnt signal/stem cell gene signature in mammary organoids.

The currently accepted view of the postnatal mammary gland is that it contains committed luminal- and basal-stem cell pools, whereas multi-potent stem cells exist during embryogenesis (Koren & Bentires-Alj, 2015; Seldin, Le Guelte, & Macara, 2017). Studies with the *waved-2* mouse model implied that strength of EGFR signals impact cell fate in the mammary gland. (Fowler et al., 1995; Sebastian et al., 1998). Pasic *et al*. had reported that EGFR signaling is critical for duct development but also that sustained EGFR signaling plays a role in myoepithelial lineage specification (Pasic et al., 2011). PI3K signals can, in principle, influence basal-versus luminal-cell fate choice comes from two exciting cancer studies. Inducible, luminal lineage-specific expression of an oncogenic allele of PI3 kinase, *PI3K^H1047R^*, can switch the cell fate from luminal to basal and vice versa, suggesting that expression of *PI3K^H1047R^* postnatal can reactivate some kind of multipotency (Koren et al., 2015; Van Keymeulen et al., 2015). In our mouse models we do not observe absolute defects in the development of either basal- or luminal-cells, even though we observe high (PI3K-)AKT signals. However, in organoids with well-controlled growth conditions generation or maintenance of the CD49f^high^/EpCAM^medium^ basal cell population that harbors stem cells is less robust. Our *Rasgrp1^-/-^* and *Rasgrp1^Anaef^* models demonstrate delayed ductal morphogenesis and reduced cellularity in the 8-week old mammary gland. The exact mammary epithelial hierarchy is unknown (reviewed in (Inman et al., 2015; Visvader & Stingl, 2014)) and it is therefore challenging at this point to pinpoint how receptor signals such as fine-tuned EGFR signals regulate the output of this hierarchy.

Our organoid assays, FACS profiles, and gene signature provide new insights how two signaling pathways can control mammary cell fate. The identity and potential of MaSC (Mammary stem cells) is a very active area of research. The ability of a single transplanted cell to reconstitute an entire ductal tree with both luminal and basal, myoepithelial cells may suggest the existence of bi-potent MaSC. In reality, this is not straightforward and is very actively debated (reviewed in (Inman et al., 2015; Visvader & Stingl, 2014)). Multicolor lineage tracing has identified K14-expressing multipotent embryonic mammary epithelial progenitors (Wuidart et al., 2018) and transplantation experiments suggested post-natal basal or MaSC multipotency (Shackleton et al., 2006; Stingl et al., 2006), but more recent work revealed that while transplanted basal cells display multipotency they remain unipotent under normal physiological conditions (Van Keymeulen et al., 2011). Other lineage tracing studies have supported MaSC unipotency and distinct stem cell pools; K8- and *Elf-5*-based tracing both established luminal-restricted progenitors (Rios, Fu, Lindeman, & Visvader, 2014; Van Keymeulen et al., 2011). However, bipotent progenitors have also been described; K5-expressing cells have been found to contribute to both mature luminal and myoepithelial lineages (Rios et al., 2014). Perhaps most indicative of a heterogeneous MaSC compartment are studies using *Lgr5* reporter mice. Lgr5 is a Wnt signaling target gene and an R-spondin receptor in intestinal stem cells (Barker et al., 2013). Reconstitution experiments with the *Lgr5* reporter have yielded contrasting results of Lgr5^+^ cells as having enriched repopulating activity compared to Lgr5^-^ cells (Plaks et al., 2013), both Lgr5^-^ and Lgr5^+^ cells having the same repopulating activity (de Visser et al., 2012; Rios et al., 2014), and Lgr5^-^ cells having enhanced activity (Wang et al., 2015). Recent single cell gene expression analyses of the human mammary gland (Lawson, Bhakta, et al., 2015; Nguyen et al., 2018; Pal et al., 2017) helps delineate and characterize rare stem and progenitor cell subpopulations. Our study here reveals that EGFR signals need to be tuned (by Rasgrp1 or other mechanisms) in order for coordinated mammary gland development to occur and that EGF can effectively dampen Wnt-induced stem cell signatures in organoid assays. Further work with well-defined growth factors in organoids combined with mouse models will be required to understand the full spectrum of signaling pathways sustaining the stem cell pool(s) in the mammary gland.

## Acknowledgments

The authors thank the members of the Roose lab and Dr. Jay Debnath for helpful suggestions and comments. Our research was supported by the Sandler Program in Basic Science (start-up), NIH-NCI Physical Science Oncology Center grant U54CA143874, NIH grant P01AI091580, a Gabrielle’s Angel Foundation grants, a UCSF ACS grant, and a UCSF Research Allocation Program (RAP) pilot grant (all to JPR), a Mark Foundation for Cancer Research Momentum Award (to PD), NCI grants R01 CA056721 (to ZW, K22 CA190511 (to D.A.L.), and R00 CA181490 (to K.K.), 1R01DK115728-01 to KCG and a fellowship NIH F31 – CA200342 (to AJS).

## Author Contributions

AS, PD, and JPR conceived the study. AS and DH performed the majority of the experiments and the analyses. PD helped initiate the project, VS helped with the transplantation assays, QHN helped with cell isolations and FACS stainings, and OG with organoid assays. JDR, WLE, AJB helped with the RNAseq experiments and data analysis. TM and KCG provided NGC Wnt factors. AS and JPR wrote the manuscript. PD, HD, KK, DL, and ZW provided insights and edited the manuscript. We dedicate this paper to our colleague and friend Zena Werb who was always open to new ideas, willing to help, and let the data do the talking.

## Materials and Methods

### Mice

Mice were handled according to the Institutional Animal Care and Use Committee regulations, described in the Roose laboratory University of California, San Francisco (UCSF) mouse protocol AN84051 ‘Ras Signal Transduction in Lymphocytes and Cancer.’ Rasgrp1 knockout (*Rasgrp1^-/-^*) mice were provided by J. Stone as described in Depeille et al. (*Nat Cell Biol*. 2015). The R519G point-mutated Rasgrp1 Anaef (*Rasgrp1^Aanef^*) mouse strain was established through ethylnitrosourea (ENU)-mediated mutagenesis of BL6 mice as previously described (Randall et al., 2009). Wild type (WT) C57BL/6 mice were bred and used as controls.

### Antibodies

Primary antibodies were obtained from the following sources and used at the indicated concentrations: P-MEK 1/2 (1:250; 2338S), P-p44/42 (1:250; 9102S), p44/42 MAPK (1:250; 9102), P-AKT S473 (1:250; 4058L), AKT (1:250; 9272), P-S6 ribosomal protein S235/236 (1:300; 2211L), S6 ribosomal protein (1:300; 2317), Cleaved caspase-3 (1:300; 9661S) from Cell Signaling; Ki-67 (1:500; ab15580), Rasgrp1 (1:100; ab37927), Cytokeratin 5 (1:100; ab52635) from Abcam; CD49f-PE-Cy7 (1:100; Invitrogen 25-0495-82); EpCAM-APC (1:63; Invitrogen 17-5791-82); CD31-450 (1:170; Invitrogen 48-0311-82); CD45-450 (1:170; Invitrogen 48-0451-82); Ter119-450 (1:170; Invitrogen 48-5921-82) from eBioscience; α-tubulin (1:2000; T6074) from Sigma-Aldrich; Troma-1 cyokertain 8 (1:100; AB_531826) from Developmental Studies Hybridoma Bank; murine Rasgrp1 m199 from Depeille et al., 2015 (Depeille et al., 2015).

Secondary antibodies were obtained from the following sources and used at the indicated concentrations: Alexa Fluor 448 (1:500 in Matrigel IF, 1:250 slide IF; A21206), Alexa Fluor 555 (1:500 in Matrigel IF, 1:250 slide IF; A21434), Alexa Fuor 568 (1:500 in Matrigel IF, 1:250 slide IF; A21069).

### Cell lines and reagents

Cell lines were cultured at 37°C in 5% CO_2_. Unmodified Eph4 cell lines were obtained courtesy of Zena Werb. Eph4 is a nontumorigeneic cell line derived from spontaneously immortalized mouse mammary epithelial cells. Eph4 cells were grown in DMEM / F21 media + 10% FBS with pen/strep. Aliquot vials of Eph4 were frozen down in growth media containing 10% DMSO. For lipofection experiments, Eph4 cells with *Rasgrp1 GFP* and *Rasgrp1 R271E GFP* plasmids were grown under selection (G418 sulfate solution, Axenia Biology). Human recombinant epidermal growth factor (hEGF) protein was purchased from Life Technologies and dissolved in PBS. Human recombinant fibroblast growth factor 2 (hFGF2) protein was purchased from Global Stem (GSR-2001) and dissolve in PBS. Insulin, transferrin, sodium selenite (ITS; 25-800-CR) was purchased from Corning. Gefitinib (S1025) and erlotinib (S1023) were purchased from Selleck Chemicals and dissolved in DMSO.

### 3D mammary epithelial colony forming cell assay

MEC clusters were first obtained following the protocol described in this manuscript. MEC clusters were then suspended in in 2mL of 0.05% trypsin/EDTA and placed in the 37°C incubator for 6 minutes. Cells were taken out every 2 minutes and pipetted up and down several times to assist in the separate of clusters into individual cells. 5 ml of HBSS + 2% fetal bovine serum was added to stop trypsinization. Now single cells were repelleted and filtered over a 100 μM strainer. MECs were counted for the colony forming assay or frozen down in DMEM / F12 + 50% FBS + 10% DMSO. For Eph4 cells, 0.25% trypsin/EDTA was used to detach cells from the plate and inspected for single cellularity. 20-μL Matrigel platforms were created in a 48-well plate and let sit for 20 minutes at 37°C to solidify. Isolated MECs are resuspended in growth factor reduced matrigel to give 2500 cells / 20μl. 20 μl of the Matrigel-cell suspension was added on top of the Matrigel platform in each well and placed in the 37°C incubator for 20 minutes to solidify. Growth factor and inhibitor medium were then added. Growth factor media used: 2.5 nM FGF, 2.5 nM EGF, 1 μM gefitinib erlotinib, in 1x pen/strep, 1x insulin transferrin, sodium selenite DMEM/F12. Size and number of colonies of organoids are followed for up to 10 days.

### Traditional Three-dimensional Mammary gland Matrigel cultures

4- to 9-week-old mice were selected for our experiments. CO_2_ euthanized mice were placed chest up and sprayed with 70% ethanol. We made a medial cut distal down the abdominal skin, which was peeled away from the peritoneum of the abdominal cavity. Inguinal (#4) fat pads were removed and cut with a scalpel until loosened. Mammary tissue and mammary epithelial cells (MECs) are transferred into 0.45 μM filtered collagenase solution; DMEM/F12 (UCSF CCF, CCFAA010-167201), 5% FBS, 50mg/mL gentamicin (Gibco, 83-50721M), 5 μg/mL insulin (Sigma, I5508), 2 mg/mL trypsin (Gibco, 27250-018) and type IV collagenase (Gibco,17104-019). Glands were shaken at 37°C for 35 min. The tissue then underwent differential centrifugation and treatment with 2 U/μl DNAse (Sigma D4263-1VL) to isolate MEC clusters. 20 μL growth-factor reduced Matrigel (BD 354230) platforms were created in glass bottom chamber slides (Lab-Tek®, 177379) or 24-well plates and incubated for 20 minutes at 37°C to solidify. Organoid density was determined before suspending MEC clusters in growth factor-reduced Matrigel at 2 organoids/μL. 20 μl of the Matrigel-cell suspension was added on top of the Matrigel platform in each well and placed in the 37°C incubator for 20 minutes to solidify. 500 μl DMEM/F12 media is added after solidification. Growth factor media used: 2.5 nM FGF, 2.5 nM EGF, in 1x pen/strep, 1x ITS, 1x DMEM/F12. Growth and branching of organoids are followed for up to 10 days. Inhibitors given at day3: EGFRi (10 μM Erlotinib), PI3K inhibition (10 μM GDC0941), or MEK inhibition (10 μM U0126). For organoids in figure 6, Rsondin2 and NGS Wnt are added at the indicated concentration.

### Single mammary epithelial cell prep for FACS

Single cell isolation for FACS from mammary tissue adopted from a protocol provided by Kessenbrock K. and Lawson D (University of California, Irvine). #1-#5 fat pads were removed from 4- to 9-week old mice and placed in a dry 10-cm dish. The tissue was chopped with a razor until slurry-like in consistency. Mammary tissue was shaken in collagenase medium (2 mg /mL collagenase type IV, Sigma C5138, in DMEM / F12, Corning 10-090) at 37°C for 1 hour. Digested tissue is spun down at 1500 rpm for 5 minutes and then washed with PBS. MECs are freed using 0.05% Trypsin/EDTA (Corning 25-0520), and excess DNA is removed using DNAse (Worthington LS002139). For FACS experiments on mammary organoids, TrypLE Select (Life Technologies 12563011) was added to wells containing organoids embedded in Matrigel pellets. Organoids were incubated at 37°C for 10 minutes. After incubation, organoid suspension was pipetted vigorously 10-15 times to further dissociate organoids to single cells. Suspension is spun down at 1500 rpm for 5 minutes and then washed with PBS. Single MECs are counted and placed into FACS tubes.

### Basal and luminal FACS stain on primary mouse MECs

We used flow cytometry to assess luminal (CD49f/EpCAM^hi^) and basal/mammary stem cell-containing CD49f^hi^/EpCAM populations (Shackleton et al., 2006; Stingl et al., 2006). Single MECs were isolated from WT, Rasgrp1^-/-^, and Rasgrp1^Anaef^ mice and placed into 500 μl DMEM / F12 FACS tubes (Falcon, 352235). Single color controls were made for CD49f, EpCAM, and Lin- (CD31, CD45, Ter119) in addition to a no stain control. FACS tubes are placed at 4°C in the dark for 20 minutes. Cells are spun down at 1500 rpm for 5 minutes and aliquoted into new FACS tubes through the cap filter. FACS tubes are kept on ice. FACS stains were performed on an LSRII and an FACS Aria II machine. Sytox blue (ThermoFisher Scientific, S3457) is added to Lin- and ALL tubes just prior to their run, in order to differentiate lin-/live (MECs) from lin+/dead (stroma).

### Mammary whole mount and carmine stain

Protocol was adapted from Kouros-Mehr H. et al. (Cell. 2006 Dec 1; 127(5): 1041–1055.) (Kouros-Mehr, Slorach, Sternlicht, & Werb, 2006). Inguinal #4 mammary fat pads were removed carefully from experimental mice. Glands were spread on glass slides (Fisherbrand, 12-550-15) and fixed overnight in 3:1 ethanol to glacial acetic acid. Slides were transferred to 70% EtOH then 50% ethanol for 10 minutes. Slides were washed with tap water slowly to remove ethanol. Slides were placed in new slide holder containing Carmine Red (carmine, Sigma C1022; aluminum potassium sulfate, Sigma A7167). Mammary tissue was then transferred to 70% then 95% ethanol for 15 minutes, and 100% ethanol for 15 x 3 minutes. Tissue was then transferred to Histo-Clear II (National Diagnostics, HS-202) for 2 hours, and then placed into fresh Histo-Clear. Slides were imaged using a dissection scope. Number and location of terminal end buds was assays. Image analysis software (ImageJ) was used to determine length of mammary duct branching.

### Cleared mammary fat pad transplantation assay for mammary epithelial organogenesis

We followed the protocol detailed in from Lawson, D.A. et al (Cold Spring Harb Protoc.; 2015(12): pdb. prot078071.). In short, mouse mammary tissue was dissociated for the isolation of 10^3^-10^6^ MECs from WT, Rasgrp1^-/-^and Rasgrp1^Anaef^ mice. MECs were resuspended in 20 μL Matrigel-GFR: primary growth medium 1:1 and kept on ice. Recipient mice aged 21-35 days were surgically cleared of their rudimental mammary epithelial duct in the #4 inguinal fat pad, located in the region of the mammary fat pad between the nipple and the proximal lymph node. MECs suspended in Matrigel mixture were then injected into the cleared mammary fat pad. Mice were followed up daily for the next week to assay for health. Epithelial organogenesis took place over 6 weeks, after which the transplanted inguinal fats pads were isolated and stained with carmine. Ductal length measured using Image J.

### Immunofluorescence of mammary whole mount sections

Mammary fat pads were dissected, placed on glass slides, fixed in 4% PFA for 1 hour, and place overnight in 30% sucrose overnight. Mammary tissue was placed in 1:1 OCT (Tissue-Tek)/30% sucrose for 1 hour. Tissue embedded in OCT. Sections (15 μm-thick, Leica CM 1950 Cryostat) were obtained, setting the cooling block to maximum cold. Sections were dried at room temperature for 1 hour, then incubated in 0.3% Triton X100 (Sigma-Aldrich) in PBS. Slides were rinsed with PBS and blocked for 1 hour (10% normal donkey serum, 3% bovine serum albumin, 0.1% Triton X100 in PBS). Primary antibodies were diluted in blocking buffer, added to the tissue, and incubated overnight at 4°C overnight.

Slides were washed with PBS and incubated in secondary antibodies (diluted in 5% normal donkey serum, 0.1% Triton X100) at room temperature in the dark for 2 hours. After additional PBS rinses, DAPI (Sigma-Aldrich) counterstain was added. Images were obtained using a motorized, upright fluorescence microscope (Zeiss Axio Imager M2, Carl Zeiss). Images were collected using Imager M2 camera. Mean corrected total cell fluorescence (CTCF) was calculated using Image J.

### Nuclear counterstain of organoids and colony-forming assay

Nuclear counterstaining for visualization was done using Hoechst® 33342 (ThermoFisher Scientific, H1399) and following the provided protocol. In short, Hoechst® dye is diluted 1:2000 in PBS and added to MECs for 10 minutes, protected from the light. Staining solution was removed and cells were washed 3x with PBS. Images were collected using an inverted fluorescence phase contrast microscope (Keyence BZ-X710, BZX Viewing Software).

### Lipofection of Eph4 MECs

Plasmids for wild-type *Rasgrp1 GFP* and catalytically dead *Rasgrp1 R271E GFP* were generated at the UCSF virus core. Eph4 cells were lipofected with *Rasgrp1 GFP* and *Rasgrp1 R271E GFP* following the Lipofectamine® Reagent (Invitrogen, 18324-012) transfection protocol. In short, 70-90% confluent Eph4 cells are incubated in media containing Lipofectamine® and DNA in Opti-MEM (Gibco, 31985-062). Eph4 cells were followed for 1-3 days and then analyzed.

### Immunofluorescence on 3D Matrigel assays

Protocol adapted from Huebner RJ et al. (Huebner, Neumann, & Ewald, 2016). Mammary epithelial cell colonies and organoids were grown in a 50:50 growth factor-reduced Matrigel: DMEM /F12 suspension. Growth factor media was removed and cells were fixed in warm 4% PFA (32% solution; 15174-3; Electron Microscope Sciences) for 25 minutes at 37°C with gentle rotation. The plate was then washed 3x in PBS at room temperature (RT) for 10 minutes. 0.5% Triton X-100 in PBS was added to wells for 30 minutes at RT. Cells were blocked for 3 hours in 10% FBS in PBS at RT. Primary antibodies were diluted in blocking buffer, placed in wells, and let sit overnight at 4°C. Wells were washed 10% FBS in PBS at RT and then blocked for 40 minutes. Secondary antibodies were diluted in blocking buffer, placed into wells, and let sit for 2 hours at RT. Wells were washed in PBS. Nuclear counterstain was added following the protocol described in this manuscript. Wells were washed with PBS and then imaged on the Keyence BZ-X710 inverted fluorescence phase contrast microscope.

### Western Blot

Cells were placed in 6-well plates and starved for 2 hours at 37°C in PBS. After resting, cells were stimulated for 0, 2, 5 and 30 minutes in EGF. Cells were lysed with ice-cold NP40 with added protease inhibitors (10 mM sodium fluoride, 2 mM sodium orthovanadate, 0.5 mM EDTA, 2 mM phenylmethylsulphonyl fluoride, 1 mM sodium molybdate, aprotinin (10 mg/mL), leupeptin (10 mg/mL), pepstatin (1 mg/mL)). After lysing for 30 minutes, lysate was centrifuged at 4°C and resuspended in 2X sample buffer. Lysates were run on pre-cast NuPAGE^TM^ 4-12% Bis-Tris gel (NP0335BOX, Invitrogen) and transferred to PVDF membranes. Protein was incubated with primary antibodies overnight. Western blots were visualized with enhanced chemo-luminescence substrate (Thermo Scientific, 32106) and imaged on a Fuji LAS 4000 image station (GE Healthcare). Bands were quantified with Multi Gauge software, and densitometry was determined within linear range of the exposure. Values were normalized to loading controls. Intensity was reported as fold difference from control sample.

### RNA Extraction and real-time PCR

Total RNA was isolated from total MECs, basal, luminal, stromal and Eph4 cells using RNeasy Kit (Qiagen). RNA underwent reverse-transcription with random primers (Invitrogen) and reverse transcriptase. RNA quantity and quality was assayed via NanoDrop spectrophotometer (Thermo Fisher) 260/60/280. SensiMix II (Bioline) was added to samples, made in triplicate and placed in real-time (RT) PCR plates (Eppendorf, 951022015) with clear film (Eppendorf, 0030132947). RT PCR was performed using the RealPlex2 (Eppendorf). Expression was normalized to ß-Actin (Mm02619580_g1, Applied Bio Systems) and quantified using the CT comparison method. This method was detailed by Eppendorf. Rasgrp1 primer (Mm00448564_m1) was obtained at Applied Bio Systems.

### RNAseq Data Analysis

STAR (Dobin et al., 2013) version 2.7.0f was used to align reads to the Mouse genome, version GRCm38.96, no adapter clipping or filtering was performed before alignment. Reads uniquely mapped to the mouse genome were used to assess expression changes between genes using DESeq2 (Love, Huber, & Anders, 2014). Variance-stabilized counts were used for creating the heatmap, while blind-stabilized counts were used for PCA. PCA was generated using the prcomp() function from the <u>stats</u> package in R. <u>Plotly</u> R package was used for making 3D PCA plots. The significance cut-off for Figure 6D was false discovery rate < 0.05. No fold change thresholding was used to restrict genes in 6D. The mean stabilized read count of each gene was subtracted from all samples’ stabilized read counts for that gene to generate color-grading, and any differences larger than the 97.5th percentile were set to the 97.5th (0.5 variance stabilized gene counts above the mean) percentile. Any stabilized gene counts differences lower than the 2.5th percentile were similarly set to the 2.5th (−0.5 variance stabilized gene counts below the mean) percentile.

### Quantification and Statistical analysis

For Western Blot analyses, densitometric intensity values were compared using paired *t*-test for pairwise comparisons.

Mammary ductal tree branch lengths were determined utilizing ImageJ analysis of captured pictures. Pixel values were transformed into metric values using provided scale information. Terminal end buds were quantified by manual counting and statistical significance was determined using paired *t*-tests for pairwise comparisons between two genotypes. Percent branched organoids were determined by inspecting 50 randomly selected in-Matrigel organoids for each mouse and averaged between the total *n* of each condition. Significance was determined by paired *t*-tests. Hoechst labelled MEC colonies were counted using captured photographs of the Matrigel pellet. A Muse^TM^ cell analyzer was used to provided quantitative cell counts. Analysis of graphs were doing using Graphpad Prism 5 software. FlowJo (v 8.8.6) was used for all FACS analyses and generation of plots.

---

**Supplemental Figure S1.**
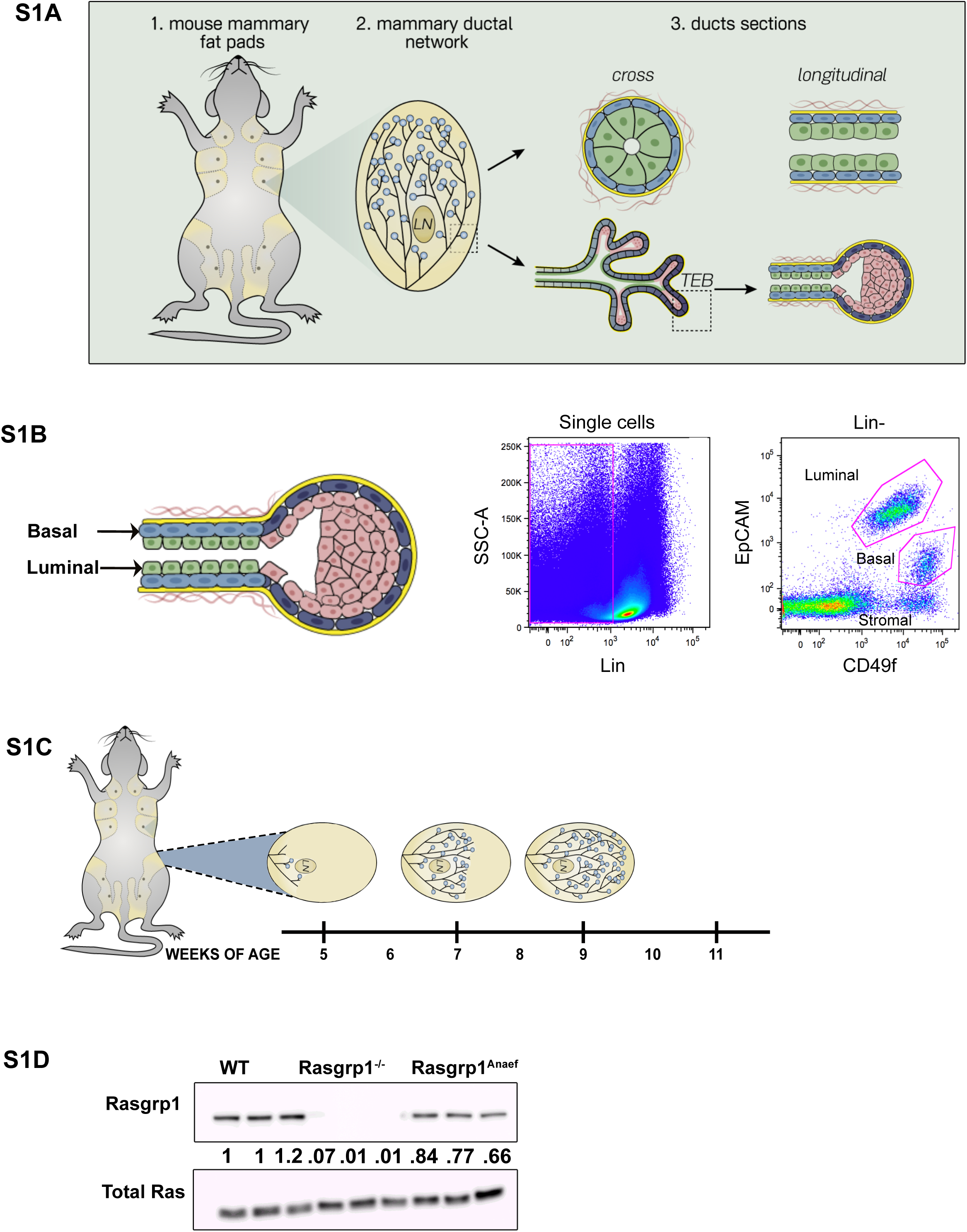
Mouse pubertal mammary gland development in C57BL/6 mice. (S1A) Cartoon depicting the architecture of the mammary ductal network and duct and TEB (terminal end bud) make-up. (S1B) Example of lineage negative (CD31^-^/CD45^-^/Ter119^-^) cells stained and analyzed for EpCAM and CD49f expression to distinguish basal and luminal cells. (S1C) Cartoon depicting pubertal mammary gland development in C57BL/6 mice. Shown is the #4 inguinal fat pad. Onset of pubertal development from the primitive duct occurs at 5 weeks, reaches an approximate midpoint at 7 weeks, and concludes at 9 weeks of age. Terminal end buds (TEBs), the driver of ductal bifurcation and elongation, subside following pubertal development. Specific time-points were selected for whole gland and mammary epithelial cell (MECs) isolation for experiments performed in the manuscript. (S1D) Western blot analysis of Rasgrp1 protein expression in murine mammary epithelial cells, normalized for cell number at extraction. WT, *Rasgrp1^-/-^* and *Rasgrp1^Anaef^* triplicate samples. Blotting for total Ras serves as loading control. Ratiometric densitometry quantification below the blot with Rasgrp1 levels arbitrarily set at 1.0 in the first WT control. n= 6 per genotype.

**Supplemental Figure S2.**
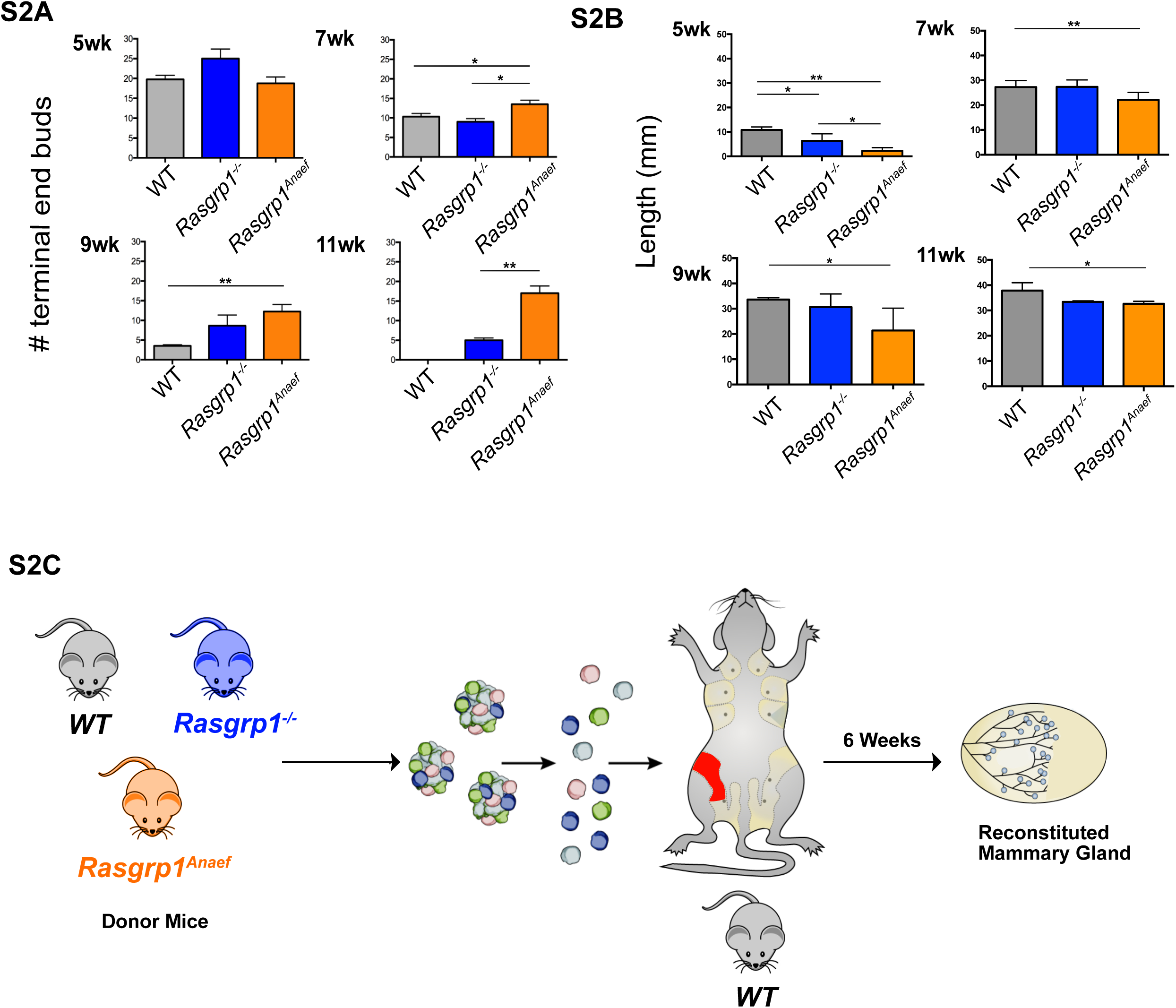
Quantification of Native mammary gland development in C57BL/6 mice and methodology for cleared fat pad transplantations. (S2A) Data accompanying Figures 2A and 2B. Mice with Rasgrp1 perturbations show elevated number of the proliferative epithelial structures, terminal end buds (TEBs). Representative images from 7-week-old, carmine-stained whole mounts. Scale bar, 2 mm. (B) Quantification of TEBs. *Rasgrp1^-/-^* and *Rasgrp1^Anaef^* mice show increased numbers of TEBs compared to their WT counterparts. Statistical significance determined by t test. (*p < 0.05, **p < 0.005; n=4). (S2B) Data accompanying Figures 2C and 2D. Rasgrp1 perturbations result in decreased ductal elongation. 7-week-old mammary whole mounts from WT, *Rasgrp1^-/-^*, and *Rasgrp1^Anaef^* were carmine stained and imaged on a stereo dissection scope. Scale bar, 10mm. (D) Quantification of ductal elongation of 5-, 7-, 9-, and 11-week old mice. Length is measured from the nipple-proximal end of the lymph node to the furthest epithelial branch. At early pubertal development, *Rasgrp1^-/-^* and *Rasgrp1^Anaef^* have significantly shorter ductal elongation (*p < 0.05, **p < 0.005; n= 4). Statistical significance determined by t test. This impairment persists through 7, 9, and 11 weeks of age. (S2C) Cartoon depicting mammary fat pad clearing and transplantation experimental design. MECs were isolated from wild type, *Rasgrp1^-/-^*, *Rasgrp1^Anaef^* mice at 7 weeks of age. MEC clusters are digested into single cells. These cells are then injected into the fat pad of a 5-week old wild type in which the endogenous primitive duct has been surgically cleared. The contralateral fat pad is left untouched, serving as a control. Transplanted mice are monitored for 6 weeks and then sacked to assess ductal outgrowth in the reconstituted mammary gland.

**Supplemental Figure S3.**
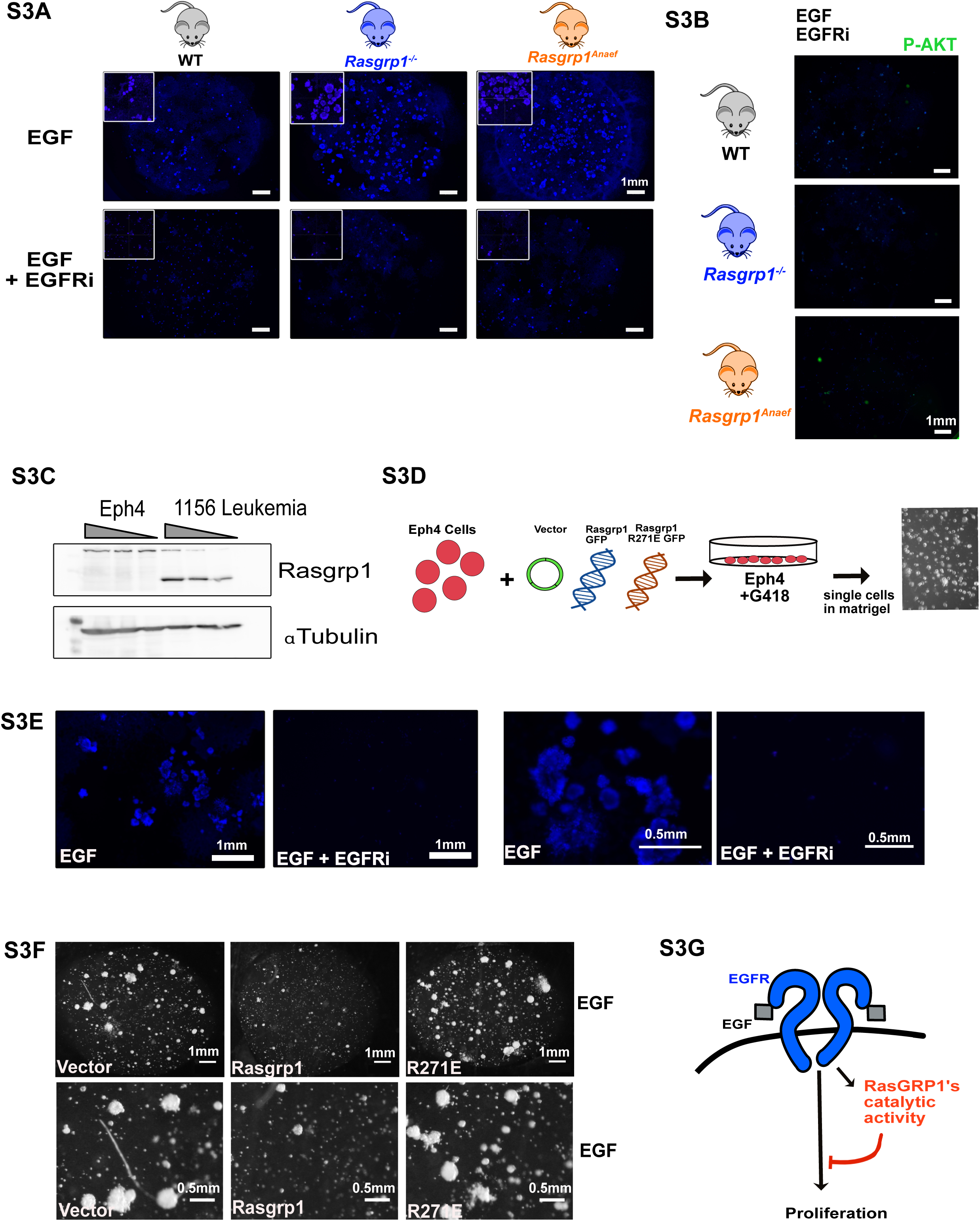
Rasgrp1 catalytic activity is required to suppress colony growth in Eph4 cells. (S3A) Data accompanying Figure 3B. *Rasgrp1^-/^*^-^ and *Rasgrp1^Anaef^* single MECs form more numerous and larger clusters in response to EGF compared to WT. In each Matrigel pellet, 2500 cells were loaded. Addition of EGFR inhibitors (EGFRi) blocks the growth of all colonies. Hoechst 33342 is used as nuclear counterstain. Scale bar, 1 mm. Zoom in images (top left). (S3B) AKT phosphorylation of colony formation assay on the three indicated, single MECs populations plated with EGFRi. Scale bar, 1 mm. (S3C) Western blot analysis for Rasgrp1 expression in the Eph4 MEC cell line and 1156 T cell leukemia cells that served as positive control. α-tubulin serves as loading control. Representative blot of 3 independent assays. (S3D) Experimental overview of Rasgrp1 expression. R271E is a mutation that disrupts the catalytic activity of Rasgrp1. (S3E) EGF stimulates robust growth of Eph4 cells in Matrigel-embedded colony forming cultures. Hoechst nuclear counterstaining was used on Eph4 colonies in Matrigel. Omitting EGF (no GF) or inclusion or EGFRi reduces colony growth. Scale bars as indicated, bottom panels are enlargements of top. (S3F) Catalytically active Rasgrp1 is required to abrogate enhanced colony formation in response to EGF. EGFRi reduces colony growth of transfected Eph4 cells. Stereo dissection scope images. Scale bars as indicated, bottom panels are enlargements of top. (S3G) Cartoon summarizing that Rasgrp1’s catalytic activity is required to inhibit EGFR-driven proliferation.

**Supplemental Figure S4.**
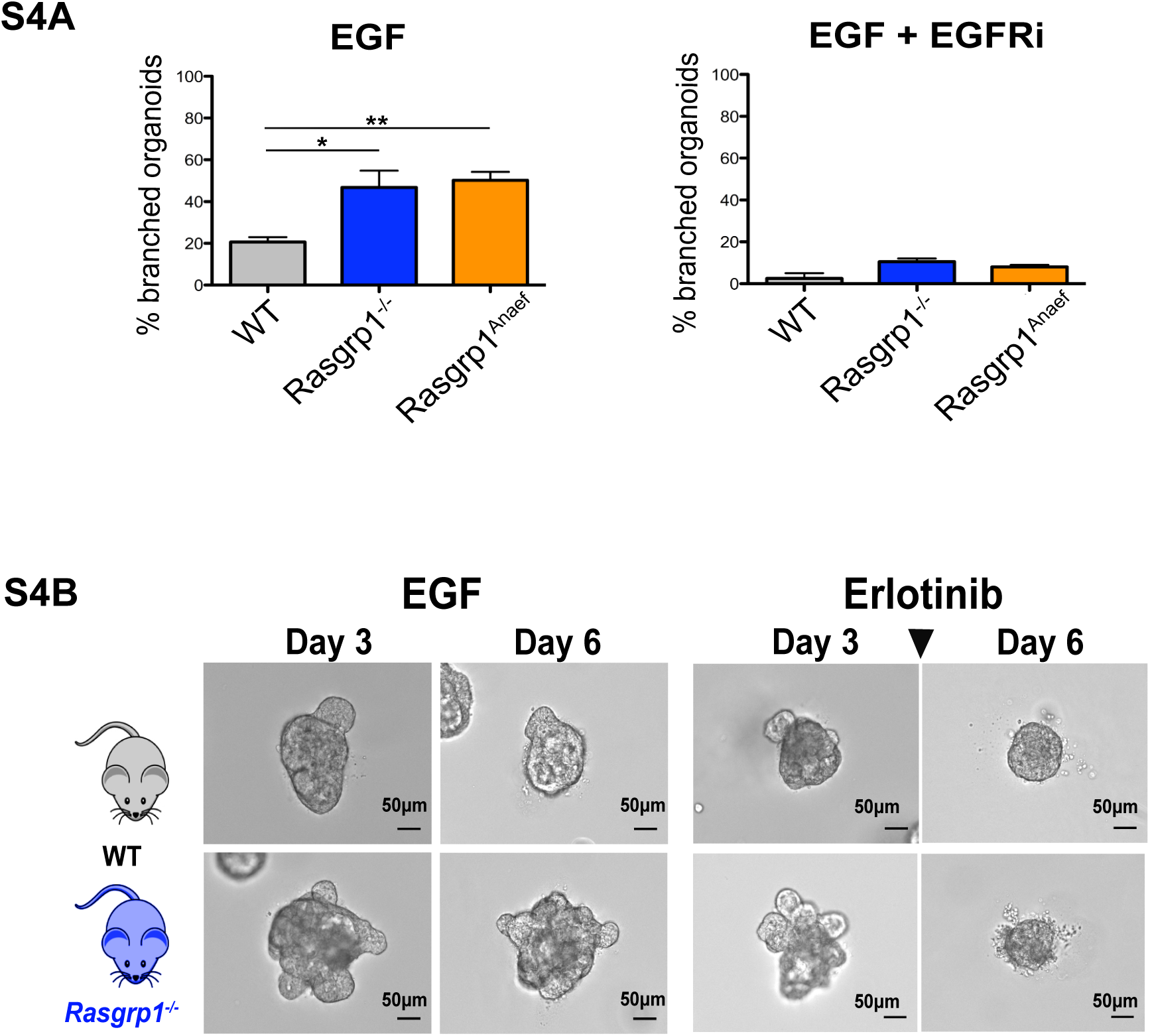
EGFR signals drive aberrant branching in *Rasgrp1^-/^*^-^ and *Rasgrp1^Anaef^*-derived 3D cultures. (S4A) *Rasgrp1^-/^*^-^ and *Rasgrp1^Anaef^*-derived 3D cultures have augmented branching in response to EGF stimulation. Assays as in Figure 4, imaging on day 6 with 3D cultures derived form 9-week-old mice. (*p < 0.05, ** p < 0.005, n=4 for each genotype and time point). Combination treatment with Erlotinib and Gefitinib EGFR inhibitors (EGFRi) reverses the gain-of-function organoid branching phenotype that occurs in EGF stimulated 3D cultures from 9-week-old *Rasgrp1^-/^*^-^ and *Rasgrp1^Anaef^* mice. No significant differences, n=4 for each genotype and time point. (S4B) Day 6 EGF-induced branching phenotypes when 3D cultures are exposed on day 3 to EGFRi. Representative results of three independent experiments. Scale bar, 50 μm.

**Supplemental Figure S5.**
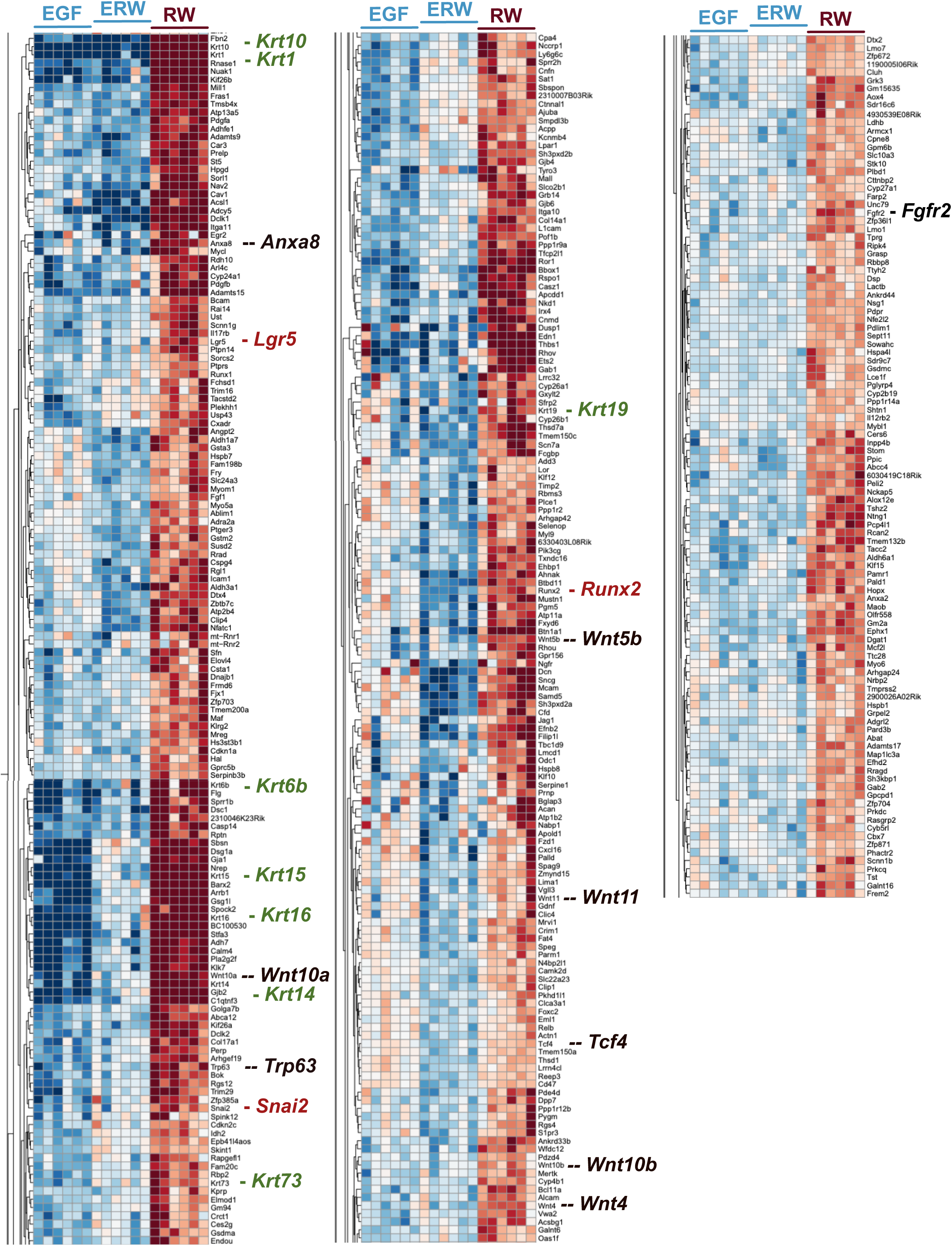
Organoid Gene expression data – Wnt and Rsponin2-driven stem cell signatures. (S5) Overview of gene expression assessed through RNAseq of organoids. Highlighted in S5 are genes that are expressed at high levels in RW medium but are expressed at lower levels in ERW or EGF.

**Supplemental Figure S6.**
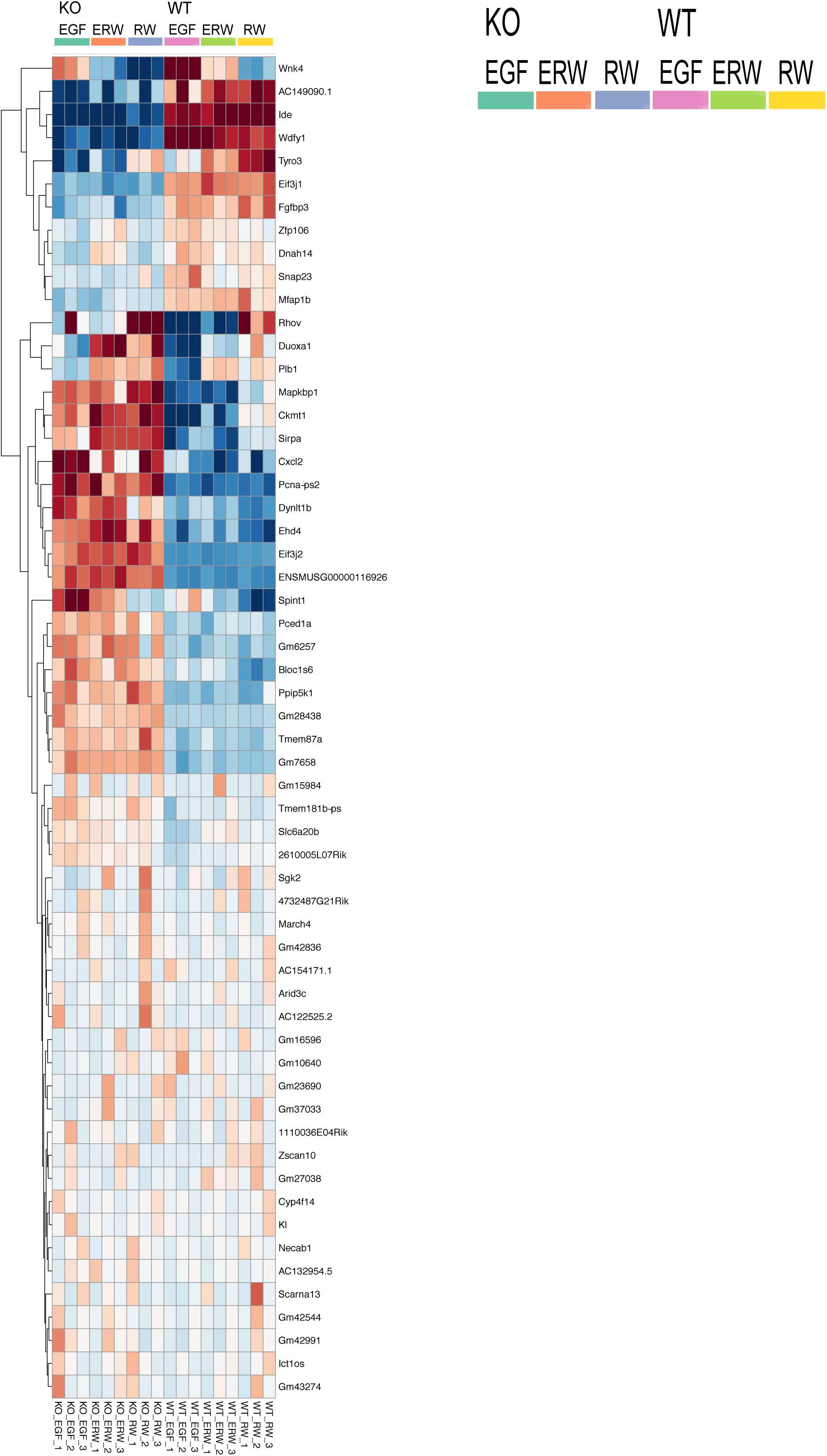
Organoid Gene expression data – Differences between WT and *Rasgrp1^-/^*^-^ organoids. (S6) Data supporting Figure 6E. DOWN in KO Wnk 4 – Serine threonine kinase AC149090.1 - Phosphatidylserine Decarboxylase Proenzyme, Mitochondrial Ide - Insulin Degrading Enzyme Wdfy1 - WD Repeat And FYVE Domain Containing 1 Tyro3 - TYRO3 Protein Tyrosine Kinase Eif3j1 - Eukaryotic Translation Initiation Factor 3 Subunit J Fgfbp3 - Fibroblast Growth Factor Binding Protein 3 Zfp106 - Zinc Finger Protein 106 Dnah14 - Dynein Axonemal Heavy Chain 14 Snap23 - Synaptosome Associated Protein 23 Mfap1b - Microfibril Associated Protein 1 UP in KO Rhov - Ras Homolog Family Member V Duoxa1 - Dual Oxidase Maturation Factor 1 Plb1 - Phospholipase B1 Mapkbp1 - Mitogen-Activated Protein Kinase Binding Protein 1 Ckmt1 - Creatine Kinase, Mitochondrial 1 Sirpa - Signal Regulatory Protein Alpha Cxcl2 - C-X-C Motif Chemokine Ligand 2 Pcna-ps2 - Proliferating Cell Nuclear Antigen Dynlt 1b - Dynein Light Chain Tctex-Type 1 Ehd4 - EH Domain Containing 4 Eif3j2 - Eukaryotic Translation Initiation Factor 3 Subunit J ENSMUSG00000116926 – pseudogene Spint1 - Serine Peptidase Inhibitor, Kunitz Type 1 Pced1a - PC-Esterase Domain Containing 1A Gm6257 – pseudogene Bloc1s6 - Biogenesis Of Lysosomal Organelles Complex 1 Subunit 6 Ppip5k1 - Diphosphoinositol Pentakisphosphate Kinase 1 Gm28438 – pseudogene Tmem87a – Transmembrane Protein 87A GM7658 - pseudogene

